# Gene regulation is commonly selected for high plasticity and low noise

**DOI:** 10.1101/2021.07.18.452581

**Authors:** Markéta Vlková, Olin K. Silander

## Abstract

Bacteria often respond to dynamically changing environments by regulating gene expression. Despite this regulation being critically important for growth and survival, little is known about how selection shapes gene regulation in natural populations. To better understand the role natural selection plays in shaping bacterial gene regulation, here we compare differences in the regulatory behaviour of naturally segregating promoter variants from *Escherichia coli* (which have been subject to natural selection) to randomly mutated promoter variants (which have never been exposed to natural selection). We quantify gene expression phenotypes (expression level, plasticity, and noise) for hundreds of promoter variants across multiple environments, and show that segregating promoter variants are enriched for mutations with minimal effects on expression level. In many promoters, we infer that there is strong selection to maintain high levels of plasticity, and direct selection to decrease or increase cell-to-cell variability in expression. Finally, taking an integrated view, we show that across all phenotypes combined, segregating promoter variants are far more phenotypically similar than would be expected given their genetic divergence. This is the consequence of both stabilizing and directional selection acting on individual phenotypes to minimize differences among segregating variants. Taken together, these results expand our knowledge of how gene regulation is affected by natural selection and highlight the power of comparing naturally segregating polymorphisms to *de novo* random mutations to quantify the action of selection.

## Main

Gene regulation plays a critical role in determining the physiology and behaviour of unicellular organisms. As such, regulation of gene expression is subject to natural selection. There are at least three aspects of gene regulation that are under selection and which affect transcription and downstream phenotypes. The first of these is expression level (the amount of protein produced from a gene). There is experimental evidence that expression levels of some genes are close to optimal (Hawkins et al., 2020; Keren et al., 2016), and furthermore, can rapidly be tuned to optimise fitness within an environment (Dekel & Alon, 2005). However, only a few studies have systematically tested the range of expression levels that have no effect on fitness, and whether this range differs between genes (Keren et al., 2016).

In contrast to the understudied role of natural selection on gene expression level, a considerable amount is known about the molecular mechanisms controlling expression level, especially in model organisms. For example, for the vast majority of genes and operons in the bacterium *E. coli*, the specific sigma factors, transcription factors, and their binding sites are well documented (de Boer et al., 2020; Ireland et al., 2020). In many cases, the specific biochemical interactions that affect expression level have been thoroughly investigated (Brewster et al., 2012, 2014; Gertz et al., 2009).

A secondary aspect of expression level is plasticity - changes in gene expression that occur when cells encounter different environmental signals. There are several aspects of phenotypic plasticity that are subject to selection; among these are the level of the transcriptional response, the speed of the response, and the sensitivity of the response (Maeda & Sano, 2006; Mangan et al., 2006; Rosenfeld et al., 2002). Although these plastic responses are well characterised, as with gene expression levels, little is known about whether stabilizing, directional, or diversifying selection is most influential in shaping phenotypic plasticity. One exception to this is the work that has been done to characterise the action of selection on the plasticity of the TDH3 promoter in yeast (Duveau et al., 2017). However, there is limited knowledge about phenotypic plasticity in bacteria and even in eukaryotes only a limited number of promoters have been studied (Duveau et al., 2017; Hill et al., 2021; López-Maury et al., 2008).

As with expression level, an abundance of research has been done on the molecular mechanisms underlying phenotypic plasticity. As noted above, many of the transcription factors that cause changes in transcription are well characterised. In addition, the specific topologies of regulatory networks that promote specific types of responses have been extensively studied (Basu et al., 2004; Becskei & Serrano, 2000; Eisen et al., 1970; Kalir et al., 2005; Mangan & Alon, 2003; Novick & Weiner, 1957; Shen-Orr et al., 2002; Smits et al., 2006), including how different topologies influence the phenotypic effects of mutations (Madan Babu et al., 2006; Mayo et al., 2006; Metzger & Wittkopp, 2019; Schaerli et al., 2018).

In contrast to expression level or plasticity, a large number of studies have quantified how selection acts on expression noise - the amount by which individual isogenic cells differ in expression level. For example, several studies have compared the noise exhibited by different promoters within the same organism to understand whether specific promoters have been selected such that they confer high or low levels of noise (Bar-Even et al., 2006; Elowitz et al., 2002; Rossi et al., 2019; Silander et al., 2012; Süel et al., 2007; Taniguchi et al., 2010; Wolf et al., 2015). There is evidence that promoters of essential genes exhibit low levels of noise, and this has been observed for both bacteria and eukaryotes (Metzger & Wittkopp, 2019; Silander et al., 2012; Urchueguía et al., 2019). In addition, modeling has shown that there can be a selective advantage for genes with high plasticity to exhibit high noise levels if the regulation of gene expression is not precise (Wolf et al., 2015). High noise levels from some promoters can lead to phenotypically distinct populations of isogenic cells coexisting in the same environment (Govers et al., 2017; Kotte et al., 2014; Ronin et al., 2017). This can be selectively advantageous as a bet-hedging phenomenon, which is sometimes beneficial in highly variable or unpredictable environments (Acar et al., 2008; Veening et al., 2008). One of the most studied examples of bet-hedging is bacterial persistence, which is known to result in a small subpopulation of cells which tolerate antibiotic exposure without acquisition of antibiotic resistance genes (Lewis, 2007).

The specific molecular mechanisms that can affect noise levels are also well-studied. For example, promoters in yeast with TATA boxes are generally more noisy than those lacking these sequences. Random mutations in TATA boxes have been found to decrease both expression noise and phenotypic plasticity (Hornung et al., 2012; Richard & Yvert, 2014). In contrast, yeast promoters lacking TATA boxes have been found to be mutationally robust (Hornung et al., 2012). In *E. coli*, specific transcription factors are associated with promoters conferring high or low levels of noise (Silander et al., 2012). Finally, there are general features of gene regulation that promote or dampen noise. For example, genes that have low rates of transcription and high rates of translation will generally have higher noise levels (Jones et al., 2014).

Here, we investigate how selection acts on different gene expression phenotypes by selecting ten *E. coli* promoters which exhibit different levels of segregating variation. We quantify expression phenotypes (expression level, plasticity, and noise) from these segregating promoter variants in three environments specific for each promoter. Finally, to elucidate whether stabilizing, directional, or diversifying selection has shaped gene regulation, we compare the expression from the segregating promoter variants, which have been exposed to natural selection, to a set of variants generated through random mutation, and which have never been exposed to selection.

## Results

### Sequence variation in segregating promoters

To characterise the relationship between genetic variation, phenotypic variation, and selection on gene regulation, we first quantified genetic variation in intergenic regions (IGRs) and open reading frames (ORFs) from 135 environmental isolates of *E. coli* (Ishii et al., 2006) that span the genetic diversity of *E. coli* (Sakoparnig et al., 2021). We assumed that a substantial portion of gene regulatory phenotypes depend on the sequence of these IGRs. We used annotations in RegulonDB to identify 605 of IGRs with known transcription start sites in *E. coli* MG1655, and which are controlled by the sigma factor σ^70 (^Santos-Zavaleta et al., 2019^)^. We also included the corresponding downstream annotated ORFs (Blattner et al., 1997). We then identified and aligned homologous regions from the 135 environmental isolates of *E. coli* (Breckell & Silander, 2020; Sakoparnig et al., 2021). To increase the likelihood of capturing homologous IGRs, we included 100 base pairs (bp) of the upstream and downstream ORFs (see **Methods**). Using these alignments, we calculated the proportion of segregating sites (PSS) and average pairwise identity (API) for each IGR (excluding the ORF portions of the alignments), as well as for the downstream ORFs (**Fig. 1a** and **Supplementary Figure 1a**). We found that the PSS in IGRs varied by more than an order of magnitude, from 0.369 (one indel and 104 SNPs across 282 bp for the IGR acting as a bidirectional promoter for the transcriptional dual regulator *melR* and α-galactosidase *melA*) to 0.007 (the IGR upstream of 30S ribosomal subunit protein *rpsM*, one SNP in 146 bp). We also observed perfect conservation for the IGRs upstream of the zinc metalloprotease *ftsH* (99 bp in length), the octaprenyltransferase *menA* (66 bp), the PF03966 family protein *ycaR* (51 bp), and the IGR that acts as as a bidirectional promoter for the transcriptional dual regulators *soxR* and *soxS* (85 bp). The API of IGRs ranged from 88.14% (one indel and 27 SNPs across 75 bp for the IGR upstream of bacterioferritin *bfr*) to 100% (*ftsH, menA, ycaR, soxR, and soxS)*.

**Figure 1:**
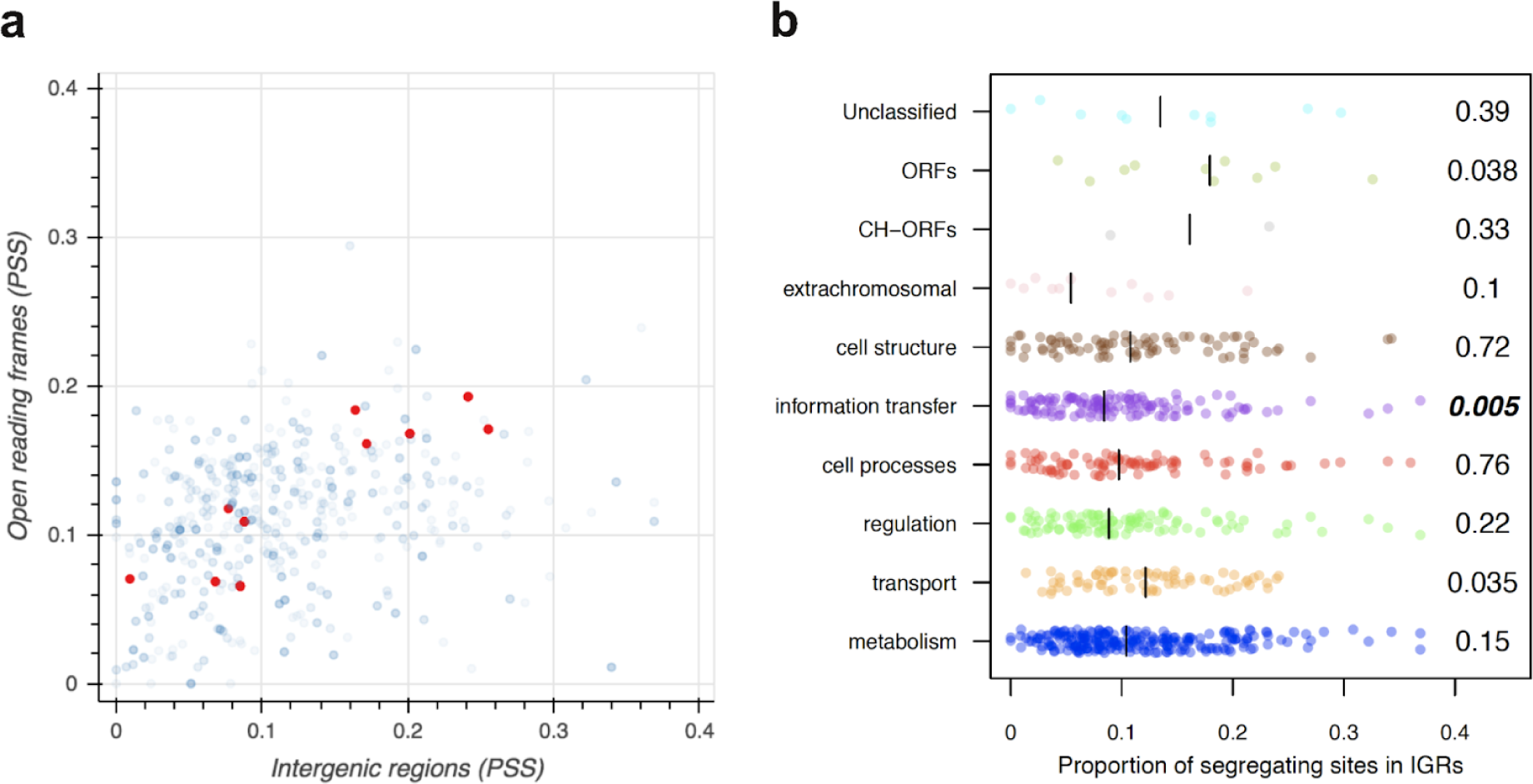
Polymorphisms in intergenic regions (IGRs) and open reading frames (ORFs) across 135 environmental isolates of *E. coli* and MG1655. a) The proportion of segregating sites (PSS) for IGRs and downstream ORFs varies by more than an order of magnitude. Each blue dot indicates PSS for an IGR-ORF pair when the IGR contains a transcriptional start site for the ORF. In red are IGRs that we selected for further study due to their different levels of sequence variation (*aldA*, *yhjX*, *lacZ, aceB*, *mtr, cdd*, *dctA*, *ptsG*, *purA*, and *tpiA*; see **Table 1**). **b**) The proportion of segregating sites for IGRs differs little among different functional groups, as classified by the downstream ORF. CH denotes conserved-hypothetical. Unclassified, ORFs, and CH-ORFs all represent groups of ORFs with very limited information on function (Serres & Riley, 2000). Numbers next to each function group represent the p-value obtained by performing the Wilcoxon rank-sum test to test for differences in PSS in each group from all other PSS values for IGRs outside of the group. Bold indicates significant p-values after the Bonferroni correction for multiple comparisons among functional groups. The black lines indicate the median PSS values of each functional group.

**Table 1:**
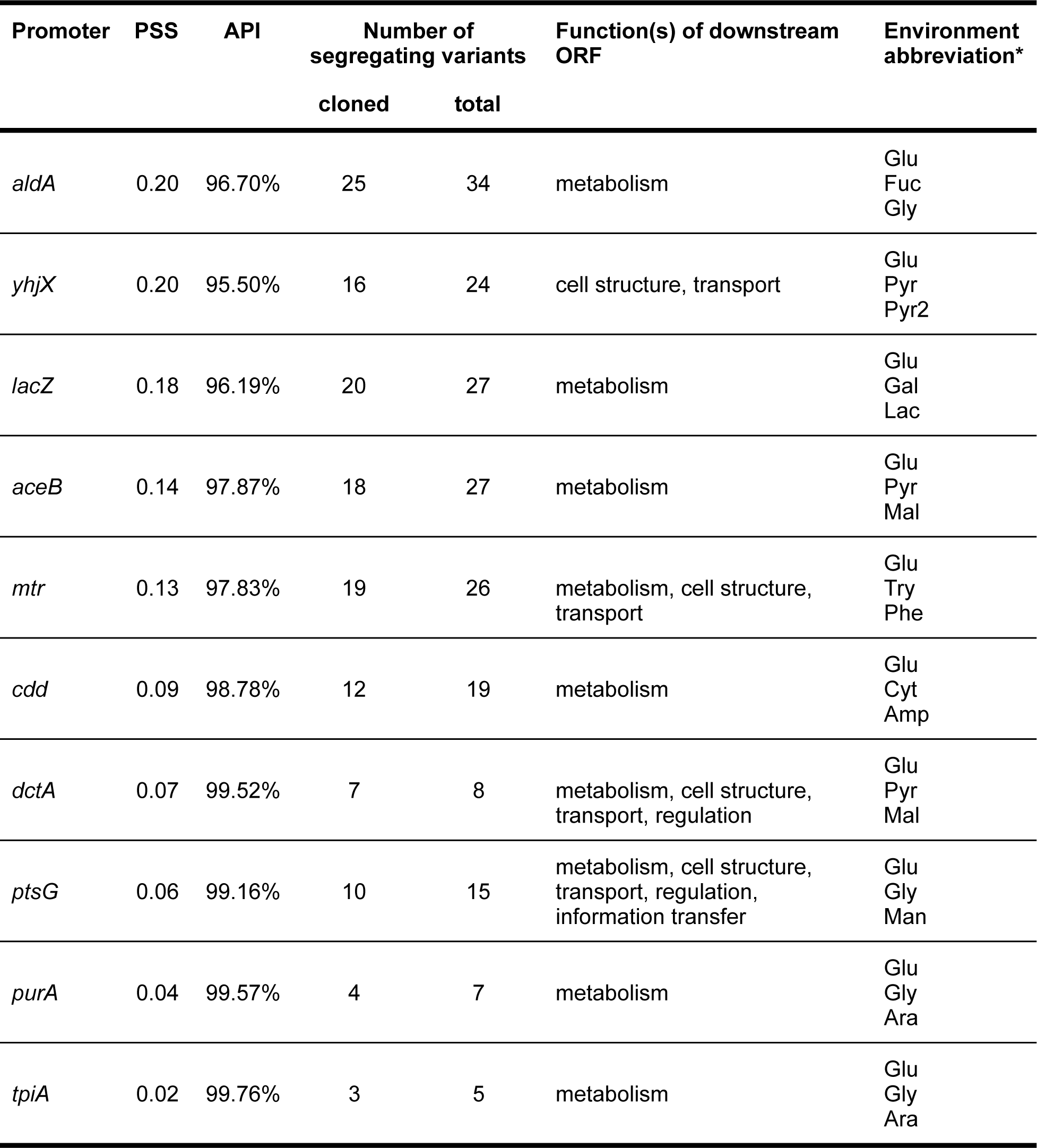
Characteristics of promoters selected for phenotypic assays. PSS - proportion of segregating sites in promoters, API - average pairwise identity in promoters. The functional groups of downstream genes were obtained using MultiFun (Serres & Riley, 2000). * For the full description of the assay environments see Table 2.

| Promoter | PSS | API | Number of segregating variants |  | Function(s) of downstream ORF | Environment abbreviation* |
| --- | --- | --- | --- | --- | --- | --- |
|  |  |  | cloned | total |  |  |
| *aldA* | 0.20 | 96.70% | 25 | 34 | metabolism | Glu<br>Fuc<br>Gly |
| *yhjX* | 0.20 | 95.50% | 16 | 24 | cell structure, transport | Glu<br>Pyr<br>Pyr2 |
| *lacZ* | 0.18 | 96.19% | 20 | 27 | metabolism | Glu<br>Gal<br>Lac |
| *aceB* | 0.14 | 97.87% | 18 | 27 | metabolism | Glu<br>Pyr<br>Mal |
| *mtr* | 0.13 | 97.83% | 19 | 26 | metabolism, cell structure, transport | Glu<br>Try<br>Phe |
| *cdd* | 0.09 | 98.78% | 12 | 19 | metabolism | Glu<br>Cyt<br>Amp |
| *dctA* | 0.07 | 99.52% | 7 | 8 | metabolism, cell structure, transport, regulation | Glu<br>Pyr<br>Mal |
| *ptsG* | 0.06 | 99.16% | 10 | 15 | metabolism, cell structure, transport, regulation, information transfer | Glu<br>Gly<br>Man |
| *purA* | 0.04 | 99.57% | 4 | 7 | metabolism | Glu<br>Gly<br>Ara |
| *tpiA* | 0.02 | 99.76% | 3 | 5 | metabolism | Glu<br>Gly<br>Ara |

In order to understand whether selection had acted in a general manner to affect the levels of polymorphism, we tested whether ORFs in certain functional categories had upstream IGRs with high or low sequence variation. We found that genes involved in information transfer were enriched for IGRs with lower levels of polymorphism (median PSS 0.084 compared to 0.104 for all other IGRs, p = 0.005; median API 98.87% compared to 98.51% for all other IGRs, p = 0.004; **Fig. 1b**, **Supplementary Figure 1b**). This lower level of variation may be due to stronger stabilising selection on the regulation of ORFs involved in information transfer. The correlation between PSS and inverted API values for IGRs is high (Spearman’s rho = 0.946, p = 4.3e-211) making the two measures interchangeable. PSS and API data for all IGRs and ORFs present in at least 130 out of 135 environmental strains are in **Supplementary File 1**.

### A system to quantify the effects of mutations on promoter phenotypes

To investigate in more detail the selective mechanisms responsible for conferring different levels of genetic variation in IGRs, we selected ten IGRs varying widely in PSS and API (**Table 1**, **Fig. 1a**, **Supplementary Figure 1a**; red points). We selected these based solely on genetic variation and for which literature data suggested phenotypic plasticity in certain environments.

For each of these ten IGRs, we cloned the segregating variants upstream of green fluorescent protein (GFP) on a low copy-number plasmid (**Fig. 2a**, **2b** and **2d**). In all cases, the cloned regions included the IGRs together with 40 - 110 bp of coding sequences flanking the IGRs to include regulatory sequences that might be present outside the IGRs. We PCR amplified the promoter region from a DNA pool of the segregating variants. Accordingly, across all ten promoters, we obtained 75% (134 out of 179) of the segregating variants (**Table 1**). We transformed all the resulting plasmids into *E. coli* MG1655, such that they were all present in a single isogenic background. We expect that if promoter phenotypes are more similar in their respective native genetic backgrounds, then this change in genetic background will only increase phenotypic differences. Thus, any conclusions we make on the similarity in the behaviour of segregating variants are conservative.

**Figure 2:**
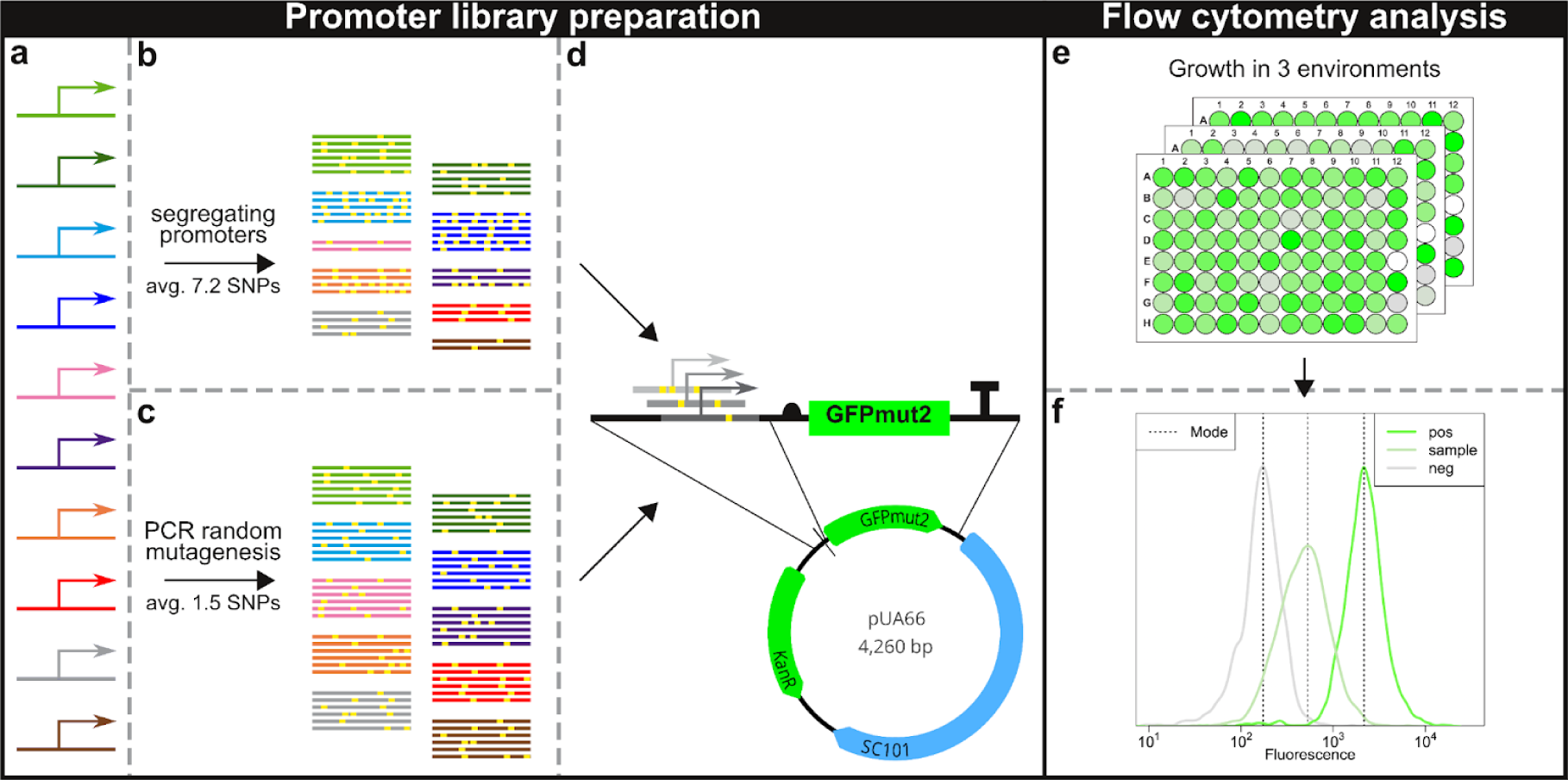
Experimental design to assay the effects of segregating and random mutations on gene expression. **a**) We isolated ten promoters (*aldA, yhjX*, *lacZ*, *aceB*, *mtr*, *cdd*, *dctA*, *ptsG*, *purA*, and *tpiA*) originating from MG1655. **b**) We then PCR amplified variants of these ten promoters segregating among environmental *E. coli* isolates from DNA pools. The average number of mutation across all segregating variants (as compared to MG1655) is 7.2 (ranging from 1 to 12.7 for individual promoters) **c**) We also performed PCR random mutagenesis using each of the ten MG1655 promoters with a target mutation rate of 1.5 mutations per promoter sequence. **d**) We cloned the resulting PCR amplicons (both segregating and random) into the pUA66 vector upstream of GFPmut2 (Zaslaver et al., 2006). We Sanger sequenced all the promoter variants to confirm the presence and location of mutations. From mutagenesis only the variants containing 1 to 3 SNPs were used for further phenotypic assays. **e**) We then cultured each of these individual promoter variants (1000 in total) in three different environments in triplicates, and **f**) quantified the modal population expression and coefficient of variation levels using flow cytometry.

We refer to these IGRs and proximal regions as promoters. We note that while differences between promoters in the expression phenotypes they confer are necessarily due to differences in the DNA sequence, it is not necessarily due to differences solely in transcription, but to any differences in the levels of transcriptional or translational regulation. Secondly, although this is a plasmid-based system, we have shown previously that expression phenotypes are well-correlated with those observed when these constructs are assayed in a chromosomal context (Silander et al., 2012).

Inclusion of the regions from the flanking ORFs had little effect on the relative genetic variabilities we observed (Spearman’s correlation for the regions inclusive and exclusive of the flanking ORFs, R = 0.918 for PSS, p = 1.09e-180; R = 0.885 for API, p = 1.05e-149; Supplementary Figure 1c **and** 1d).

For each promoter we then identified specific environments in which it exhibited different expression levels. As it is well-established that glucose is the preferred carbon source for *E. coli* (Monod, 1949), we included this as an environment for all promoters, assaying the activity during exponential growth in M9 minimal salts media with 0.4% glucose (Glu, **Table 1** and **2**). We used information from the literature on the behaviour of the MG1655 promoter variants to identify additional environments in which we expected a promoter to exhibit different expression levels. We confirmed this behaviour using flow cytometry. In this way, for each promoter, we identified two additional environments in which we observed differential expression (**Table 1** and **2**, **Figure 2e** and **2f**). We found that all segregating variants behaved consistently in all the environments that we tested. In the case of *yhjX,* we were not able to obtain expression levels higher than cellular background fluorescence in any environment that we tested except pyruvic acid. Nevertheless, we found that the promoter was very responsive to the concentration of pyruvic acid. We thus used two different concentrations of pyruvic acid as a carbon source to achieve differential levels of expression.

### Relationship between segregating genetic variation and phenotypic variation is correlated

We hypothesized that the differences we observed in the genetic variation present for different promoters might be a consequence of differences in the range of optimal expression levels. If the optimal range was very narrow, then all variants of that promoter should exhibit similar expression levels, and new mutations that affect expression should be filtered by selection, decreasing genetic diversity. In contrast, if there were a wide range of optimal expression levels within an environment, then variants can differ in expression levels, selection would act weakly against new mutations, and segregating genetic diversity would be high. Under this hypothesis, there should be a positive correlation between genetic variation and phenotypic variation. A second possibility is that segregating variants have very little effect on phenotype and are selectively neutral. In this case we would expect no correlation between genetic variation and phenotypic variation.

To test for such a correlation, for each promoter, we measured the modal population expression level from each segregating variant in each environment. In all cases, we measured the modal population expression from three full biological replicates (**Methods**). As a measure of phenotypic variation, we calculated the standard deviation in log expression level across all segregating variants of each promoter. We found that promoters with more segregating genetic variation tended to have larger phenotypic variation in expression (**Fig. 3a** and **3b, Supplementary Figure 2a** and **2b**; Spearman’s rho = 0.666, p = 5.8e-05). While this evidence does not conclusively show that the lower level of genetic variation in some promoters is due to the increased strength of stabilising selection, it shows that our experimental system is capable of detecting the effects of segregating mutations on expression phenotype.

**Figure 3:**
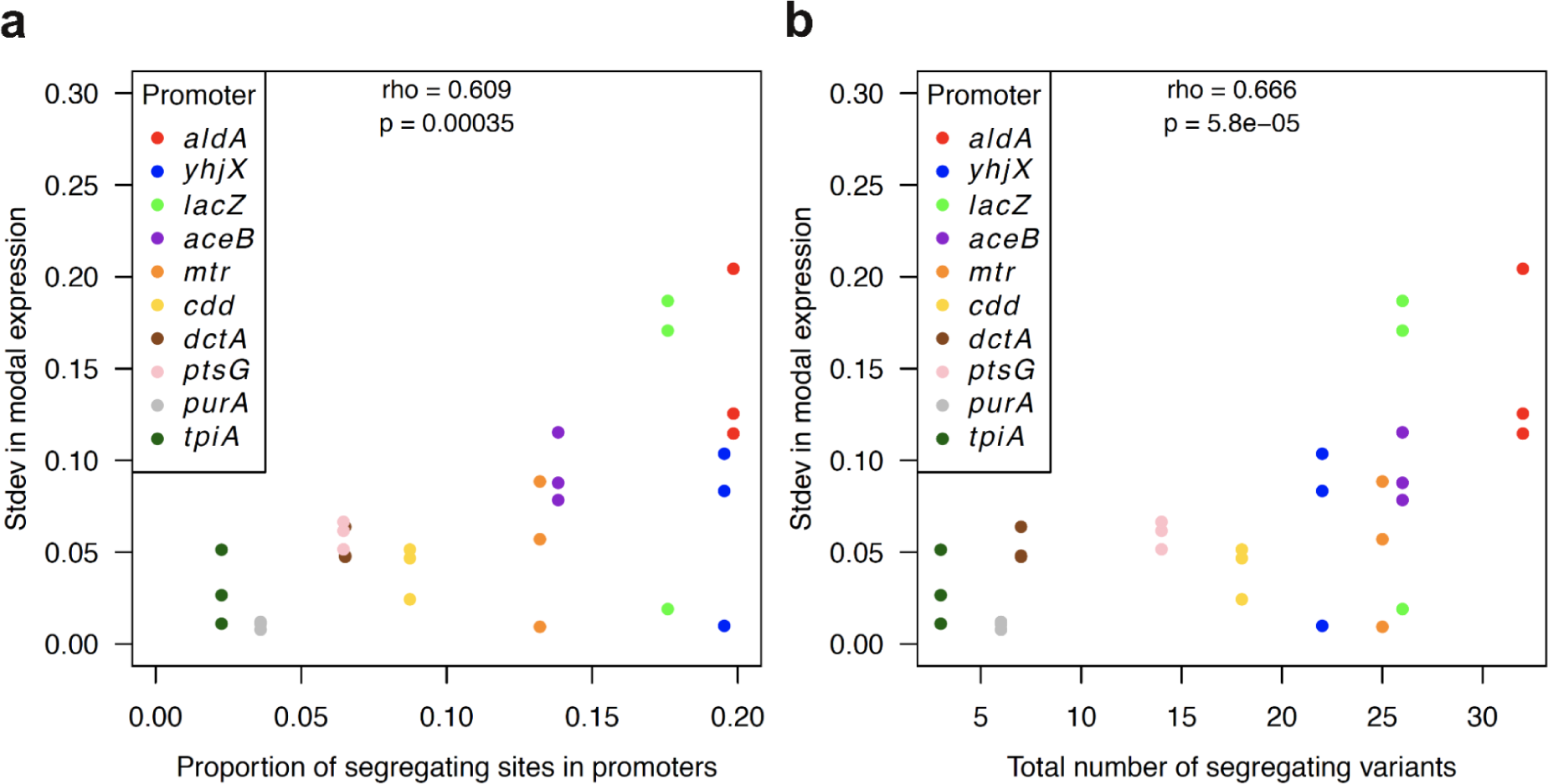
Segregating genetic variation in promoters correlates with phenotypic variation in expression levels. **a**) and **b**) Standard deviations in modal expression levels from segregating variants are correlated with the genetic variation of the promoter (IGR with 100 bp flanking regions). Panel **a** shows the correlation with PSS and panel **b** shows the correlation with the total number of segregating promoter variants. For each promoter, the standard deviation of modal population expression was measured in three environments (three dots per promoter, **Table 2**). The rho and p-values were calculated using Spearman’s correlation test.

**Table 2:** Promoter-environment combinations. All environments included M9 minimal salt media supplemented with MgSO_4_, CaCl_2_, and 50µg/ml Kanamycin.

| Promoter | Assay media | Incubation time prior flow cytometry | Environment abbreviation |
| --- | --- | --- | --- |
| *aldA* | 0.4% glucose | 4.5h | Glu |
|  | 0.4% fucose | 4.5h | Fuc |
|  | 0.4% glycerol | 4.5h | Gly |
| *yhjX* | 0.4% glucose | 4.5h | Glu |
|  | 0.4% pyruvic acid | 4.5h | Pyr |
|  | 0.2% pyruvic acid | 4.5h | Pyr2 |
| *lacZ* | 0.4% glucose | 4.5h | Glu |
|  | 0.4% galactose | 4.5h | Gal |
|  | 0.4% lactose | 4.5h | Lac |
| *aceB* | 0.4% glucose | 4.5h | Glu |
|  | 0.4% pyruvic acid | 4.5h | Pyr |
|  | 0.4% L-malic acid | 7.0h | Mal |
| *mtr* | 0.4% glucose | 4.5h | Glu |
|  | 0.4% glucose + 1mM L-tryptophan | 4.5h | Try |
|  | 0.4% glucose + 1mM L-phenylalanine | 4.5h | Phe |
| *cdd* | 0.4% glucose | 4.5h | Glu |
|  | 0.4% glycerol + 2mM Cytidine | 4.5h | Cyt |
|  | 0.4% glycerol + 3mM Adenosine monophosphate | 4.5h | Amp |
| *dctA* | 0.4% glucose | 4.5h | Glu |
|  | 0.4% pyruvic acid | 4.5h | Pyr |
|  | 0.4% L-malic acid | 7.0h | Mal |
| *ptsG* | 0.4% glucose | 4.5h | Glu |
|  | 0.4% glycerol | 4.5h | Gly |
|  | 0.4% D-mannose | 4.5h | Man |
| *purA* | 0.4% glucose | 4.5h | Glu |
|  | 0.4% glycerol | 4.5h | Gly |
|  | 0.4% L-arabinose | 5.5h | Ara |
| *tpiA* | 0.4% glucose | 4.5h | Glu |
|  | 0.4% glycerol | 4.5h | Gly |
|  | 0.4% L-arabinose | 5.5h | Ara |

### Effects on expression level are environment-dependent

Segregating promoter variants have evolved under the pressure of natural selection. Despite this, some segregating variants differ by up to 27 SNPs from the MG1655 variant (the putative transporter *yhjX*), and we have found that these mutations have detectable phenotypic effects - promoters with larger levels of sequence variation tend to have a larger range in modal expression levels (**Fig. 3a** and **3b**). To investigate how selection has affected the level of segregating variation, we created a set of promoter variants with mutations that had not been subject to selection. To do this we used PCR mutagenesis on the MG1655 promoter variant, as a representative of segregating versions of each of the ten promoters. We cloned these upstream of GFPmut2 gene on a plasmid and transformed them into MG1655, creating a set of promoter variants exactly analogous to the segregating variants, but containing mutations that had never been subject to the action of selection (**Fig. 2c** and **2d**).

To identify the specific mutations that each of these new promoters contained, we sequenced 192 clones for each of the ten promoters (1,920 clones in total; see **Methods**). We filtered out all duplicated variants and any containing none or more than three *de novo* random mutations. Next, for nine of the ten promoters, we selected 93 randomly mutated variants to produce a well-characterised library of promoter variants containing mutations that have never experienced the action of natural selection. For the tenth promoter, *lacZ*, we obtained only 29 random variants (see **Methods**). Given the small sample size, selective effects will be more difficult to detect, as statistical significance will generally require larger effect sizes.

Using these libraries, we first characterised the effects of the randomly introduced SNPs on expression level, considering only promoters with a single SNP (**Fig. 4a**). As expected (Belliveau et al., 2018; Kinney & McCandlish, 2019), we found that mutations with the largest effects on expression in all environments and promoters were present in the regions required for σ^70^ binding (-35 or -10 elements) or annotated transcription factor (TF) binding sites (p = 2.27e-09 for larger fold-changes in expression for SNPs within vs. outside of σ^70^ and TF binding sites, Wilcoxon rank-sum test). However, these data also showed that both the direction (i.e. an increase or decrease in expression level) and size of the mutational effects were dependent on the growth environment. In some promoter-environment combinations almost half of the mutations resulted in increased expression, while in a different environment these same mutations decreased expression. For example, comparing the effects on expression level in pyruvic acid vs. L-malic acid, 44.2% of all single mutations (both inside and outside TF binding sites) in *aceB* and 31.11% of all single mutations in *dctA* had opposing effects (**Fig. 4b**). In other cases, the effect sizes of the mutations were environment-dependent (e.g. compare *aldA* in glucose vs. fucose or *yhjX* in glucose vs. pyruvic acid; **Fig. 4a**).

**Figure 4:**
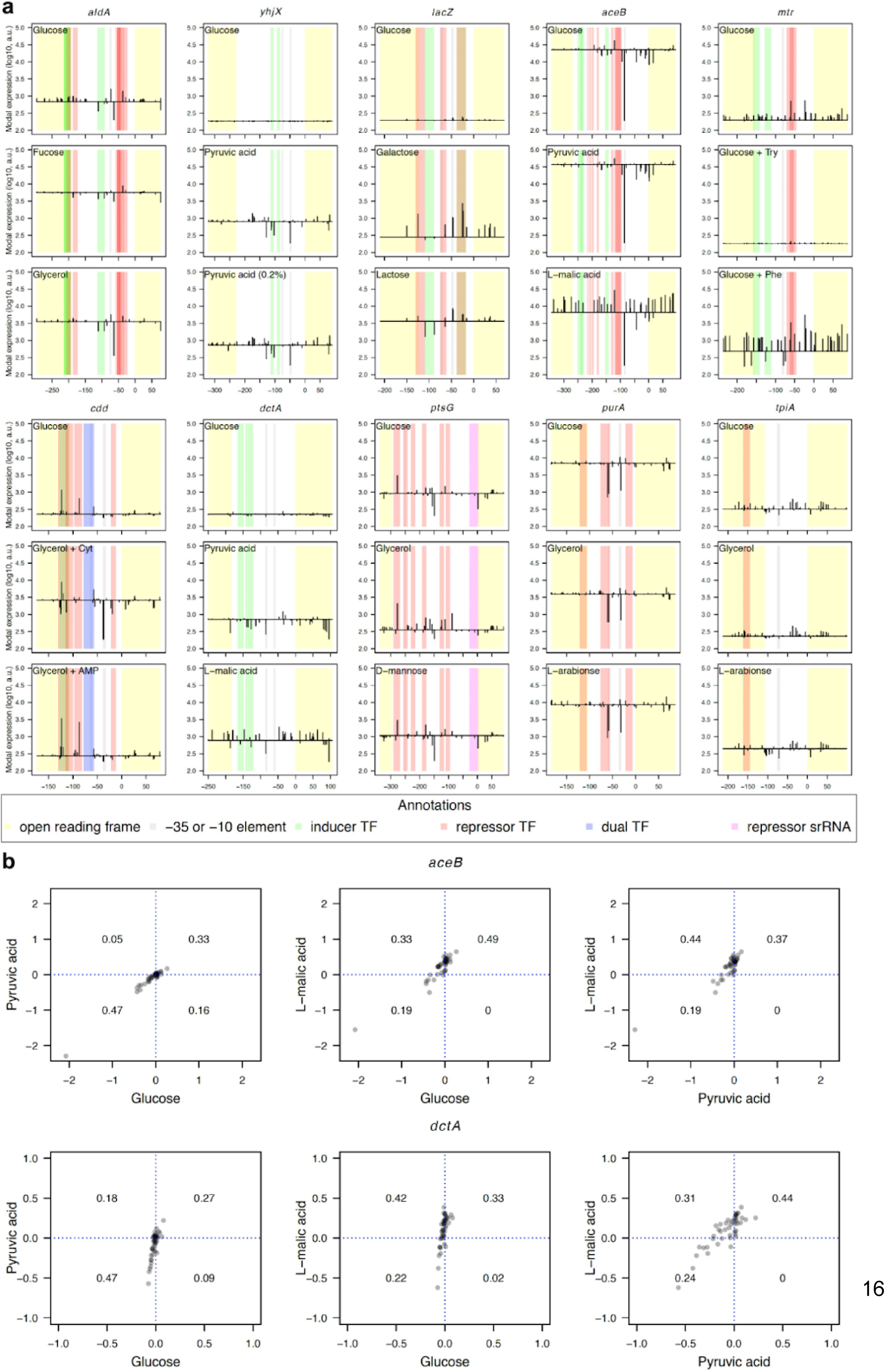
The effects of random mutations on expression level are promoter and environment dependent. **a**) Changes in expression from MG1655 promoter variants due to single random mutations for all environments. For each promoter, the x-axes indicate the position relative to the start codon of the gene downstream of each promoter. The horizontal black line indicates the expression level of the MG1655 promoter variant in that environment. The vertical black lines show the direction and size of the change in expression level when the MG1655 promoter variant is mutated at that specific position. While in certain environments some mutations cause no change in expression phenotype in other environments, the same mutations can cause up to 10-fold changes in expression. The results shown here are only from random promoter variants containing a single mutation relative to the MG1655 promoter variant. Note that for *mtr*, the MG1655 variant has a mutation in the downstream GFP, causing lower fluorescence (**Supplementary Note**), and thus the random mutations exhibit larger effects when the promoter is activated. The brown stripes result from overlapping repressor and activator TF binding sites. **b**) Comparison of differences in modal expression from random variants with single SNP relative to the MG1655 variant for *aceB* and *dctA* promoters. Differences are always compared between a pair of environments. Blue dotted lines indicate a null change in expression as compared to the MG1655 variant. These lines also divide each plot into four quadrants. Data points in the bottom left and top right quadrants show changes in expression in the same direction (decrease or increase) in both environments. Variants clustering in top left and bottom right quadrants exhibit changes in the opposite directions in the two environments (increase in one environment and decrease in the other). The numbers in each quadrant represent the proportion of variants falling into that quadrant. Each data point in **a** and **b** is a result of three full biological replicates (**Methods**).

We note that the MG1655 *mtr* variant is expressed at lower levels, which is likely caused by a non-synonymous mutation in GFP (**Supplementary Note**). The changes in expression in **Fig. 4** for the *mtr* promoter, most notably for glucose with phenylalanine (Glucose + Phe) thus do not accurately represent the magnitude of the changes to native MG1655 promoter variant. All together, these results emphasize that in order to gain general insights into the effects of mutations on gene expression phenotypes, it is necessary to measure promoter activity across different growth environments. Not only did we observe that the relative effects of random mutations on promoter activity varied across promoters, but both the size and direction of effects varied across environments (**Fig. 4b**). However, due to the action of selection, this may not be true for naturally segregating promoter variants - for example all large-effect mutations may be filtered by selection. To quantify the action of selection, we next investigated differences between the phenotypic effects of naturally segregating and *de novo* random mutations.

### Segregating polymorphisms are enriched for mutations with small effects on expression level

We first compared the expression levels of each of the 93 random variants per promoter (29 in the case of *lacZ*) to the expression levels of segregating promoter variants. Specifically, for each promoter and each environment, we tested whether expression levels among random mutants varied more than expression levels among segregating promoter variants. In all cases in which we observed significant differences, we found that random promoter variants differed more in expression levels than did segregating variants (**Fig. 5**). This is despite the far larger numbers of segregating mutations (on average 7.2 mutations different compared to MG1655) compared to randomly generated mutations (on average 1.5 mutations different compared to MG1655). This shows conclusively that segregating mutations are enriched for mutations with small effects on expression levels, across a range of environments and promoters having different levels of segregating diversity (**Fig. 5**).

**Figure 5:**
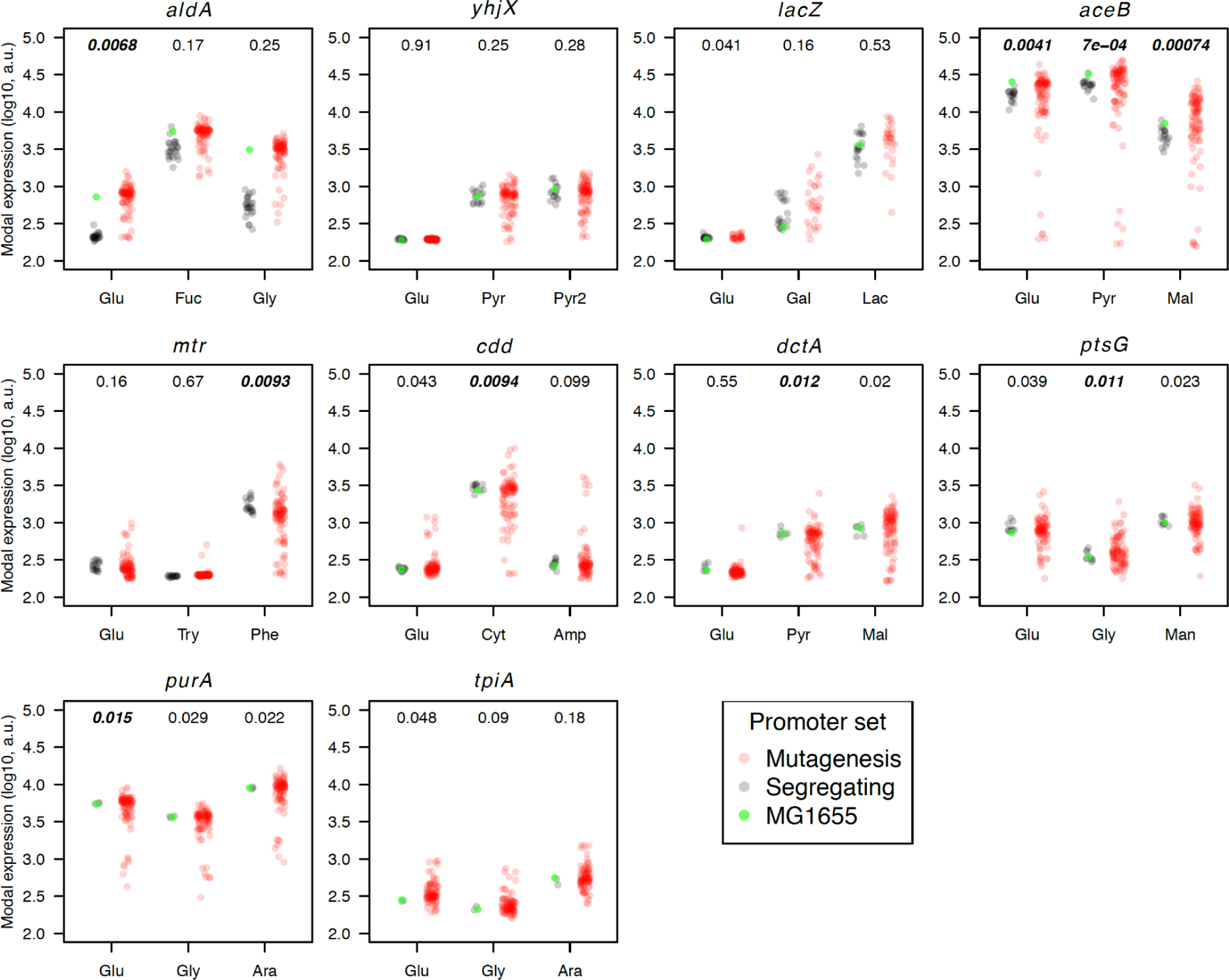
Selection acts against mutations with large effects on expression levels. We calculated the modal population expression level for all segregating and random promoter variants, and tested whether segregating variants differed in expression levels more or less than random variants. If segregating promoter variants exhibit similar expression levels (i.e. they have low variance), compared to random promoter variants, then we can conclude that large-effect mutations are filtered out via selection. For each promoter, the expression level of each group of variants is plotted. Segregating variants are shown in grey, random variants in red, and the MG1655 variant in green. The expression levels during growth in each of three different environments are shown in each plot and indicated on the x-axis (**Table 2**). Here, as in the other figures, the panels are arranged in decreasing order of segregating genetic variation (PSS). As noted in Fig. 4, the MG1655 *mtr* promoter variant has a mutation in GFP, causing lower fluorescence, which is most apparent when the promoter is activated (**Supplementary Note**), we thus omitted it from the calculations. The numbers above each pair of segregating and random variants is the p-value obtained through the Fligner-Killeen test to test for differences in variances between the two groups. The numbers in bold indicate significant p-values after the Bonferroni correction for multiple comparisons within the promoter.

In the majority of cases, the MG1655 promoter variant is representative of other segregating versions. However, we found two cases in which the MG1655 promoter variant behaved differently than all other segregating promoter variants. In the case of *aldA*, all segregating variants exhibited lower expression levels than the MG1655 variant across all conditions, except for a single variant in fucose (**Fig. 5**). In addition, the expression levels of promoter variants obtained via random mutagenesis of the MG1655 variant clustered around MG1655. The increased expression in the MG1655 variant is likely due to a single SNP that changes the -35 element from TGCCGT to TTCCGT, which is more similar to the canonical -35 motif (Harley & Reynolds, 1987). In addition, the MG1655 *mtr* variant was expressed at lower levels likely linked to a single SNP in the GFP gene, as mentioned above (**Supplementary Note**) and was thus excluded from the analysis.

### Segregating polymorphisms are selected to maintain high levels of phenotypic plasticity

A key role of promoters is to control expression of downstream genes in order to provide optimal levels of gene product in particular environments, i.e., phenotypic plasticity. To understand how selection acts on phenotypic plasticity, we again compared the phenotypes of the library of random variants to the segregating variants. We hypothesised that frequently, genes involved in the metabolism of specific substrates will be selected such that their expression is highly plastic (i.e. environment-dependent). We thus compared the environmental dependence of expression levels for all segregating and random promoter variants in all three environments. When considering two environments, highly plastic promoters should result in high expression levels in one environment, and low expression in the second environment. Less plastic promoters would have similar expression levels across both environments. We quantify this plasticity by calculating the distance of each promoter variant from an isocline specifying identical expression in both growth environments (**Fig 6a** and **Supplementary Figure 3**). This same logic can be applied to three environments: promoters with similar levels in all three environments are defined as having low plasticity, while strong differences in expression levels in each of the three environments will be observed in highly plastic promoters. Thus to quantify plasticity, we calculated the Euclidean distance of each promoter variant in three dimensions (i.e. the three environments) from an isocline representing equal expression levels in all three environments (**Fig. 6b**). This distance is 0 for promoter variants that are expressed at the same level in all environments, and greater than zero for promoter variants that are expressed at different levels, with a maximal value that is dependent on the maximum difference in expression levels between any two environments.

**Figure 6:**
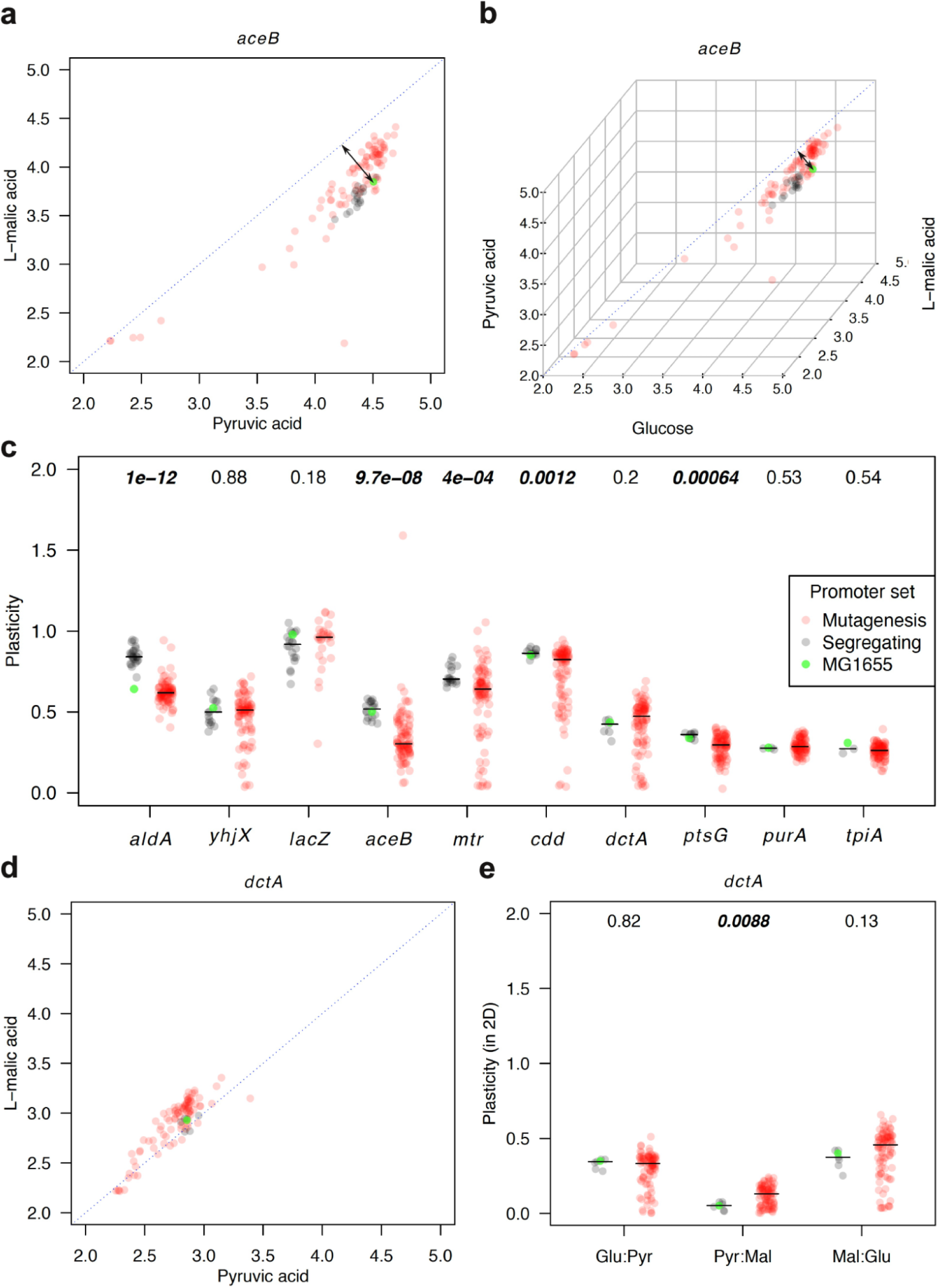
Selection on plasticity. Comparison of expression levels from promoters in a combination of environments. **a**) For the *aceB* promoter in pyruvic acid and L-malic acid, the segregating variants tend to have high expression in pyruvic acid and low expression in L-malic acid, indicating high plasticity in these two dimensions. **b**) The same projection as **a** but using all three environments. The blue dotted lines in **a** and **b** indicate the isocline of equal expression levels in all environments, i.e, no phenotypic plasticity. The black arrows illustrate the plasticity for the MG1655 promoter variant (measured as the minimal distance of each point from the blue isocline, i.e., x = y for **a** and x = y = z for **b**). The further from this isocline a promoter variant is, the higher its phenotypic plasticity. **c**) Plasticity calculated as in **b** for all environments and all ten promoters. Segregating promoter variants often have higher levels of plasticity across the three environments. The promoters are arranged in decreasing order of segregating genetic variation (PSS). The MG1655 variant of the *mtr* promoter was omitted from calculation due to a SNP in GFP (**Supplementary Note**). **d**) Expression of all *dctA* variants in pyruvic acid and L-malic acid. In this promoter, we detected low levels of plasticity in segregating variants as compared to random variants in these two environments. **e**) Calculating the plasticity of *dctA* using pairs of environments we found that in pyruvic acid and L-malic acid segregating promoters exhibited lower plasticity (p = 0.009). Numbers above each pair of segregating and random variants in **c** and **e** are the p-values obtained via Wilcoxon rank-sum test to test for differences between the two groups. The numbers in bold indicate significant p-values. The horizontal black lines in **c** and **e** indicate the median plasticity values of each group.

As shown above (**Fig. 5**), many random mutations have large effects on expression, and we observed many mutations that increased as well as decreased expression levels. However, this was not true for plasticity. Only rarely did random mutations increase plasticity. For four out of the ten promoters, we found significant differences in plasticity between segregating and random variants (**Fig. 6c**; excluding *aldA* promoter results - see below), with random promoter variants exhibiting lower plasticity. Using the median plasticity of segregating promoter variants, 92.47% (*aceB*), 75.27% (*mtr*), 76.34% (*cdd*), and 86.02% (*ptsG*) of the random variants exhibited lower plasticity. This strongly suggests that segregating promoter variants are under strong selection for high plasticity in these cases. Furthermore, this is almost certainly a conservative estimate of the strength of selection on plasticity, as we have assayed expression in only three environments.

Interestingly, when we calculated plasticity only for pairs of environments (**Supplementary Figure 3**) we found a case when the random variants exhibited significantly higher plasticity than the segregating variants - *dctA* in pyruvic vs. L-malic acid (76.34% of random variants have higher plasticity than is the median for segregating variants; median plasticity of 0.184 for random variants as compared to a median of 0.073 for segregating variants, p = 0.009; **Fig. 6d** and **6e**, Pyr:Mal). However, the difference in plasticity across all three environments was not significant (p = 0.199; **Fig. 6c**). This was the only case in which the data implied that there was selection for low plasticity, i.e., for similar levels of expression across different environments. Finally, in the case of *aldA*, we found that the MG1655 variant exhibited considerably lower plasticity than all other *aldA* segregating variants (**Fig. 5**). The random variants have similar plasticity as MG1655, resulting in a strong difference in plasticity between the segregating and random variants. (**Fig. 6c**, *aldA* and **Supplementary Figure 3**, *aldA*). Despite this, if we compare the plasticity of the random variants to that of MG1655, we find that 67.74% exhibit lower plasticity. This suggests that even though the MG1655 variant has lower plasticity than other segregating variants, selection has still acted to filter variants that decrease plasticity.

### Segregating polymorphisms are enriched for mutations that both increase or decrease noise

Finally we also looked at the role of selection on noise levels among our ten promoters. Here, we use the term “noise” to refer to the differences between isogenic cells in their protein expression level. Promoters conferring high levels of noise result in isogenic cells having different protein expression levels, while promoters conferring low noise result in almost all cells having the same expression level. We note that noise is highly dependent on expression level due to the stochastic production of mRNA and protein. In general, promoters that confer low expression exhibit higher noise, while promoters that confer high expression exhibit lower noise. Because there is strong selection on the expression level of a protein, we first calculated a metric that allowed us to decouple noise from expression level. Specifically, we calculated the vertical deviation from a fitted smooth spline on the log of the modal expression levels versus the coefficients of variation (**Methods**). We then fitted the spline using all variants (segregating and random) for a particular environment (**Supplementary Figure 4**). Thus, positive deviations from the fitted spline indicate promoters with high noise, while negative values indicate promoters conferring low noise (Silander et al., 2012).

We found that eight out of ten promoters had significantly different levels of noise (after Bonferroni correction for multiple tests for each promoter) in segregating variants as compared to random variants in at least one environment (**Fig. 7**). In the majority of cases, noise was lower for segregating variants. As we calculated the noise metric separately for each promoter and each condition, this strongly suggests that selection has acted directly to decrease noise. The differences in noise are not due to differences in regulatory inputs, or to growth conditions, but to subtle changes in the sequence of the promoter and the resulting phenotypes. It is possible that some of these effects are mediated by changes in the binding strength of various transcription factors. However, this is unlikely to be the primary factor. We would expect the major effect of these changes to be expression level, and this noise metric explicitly accounts for expression level.

**Figure 7:**
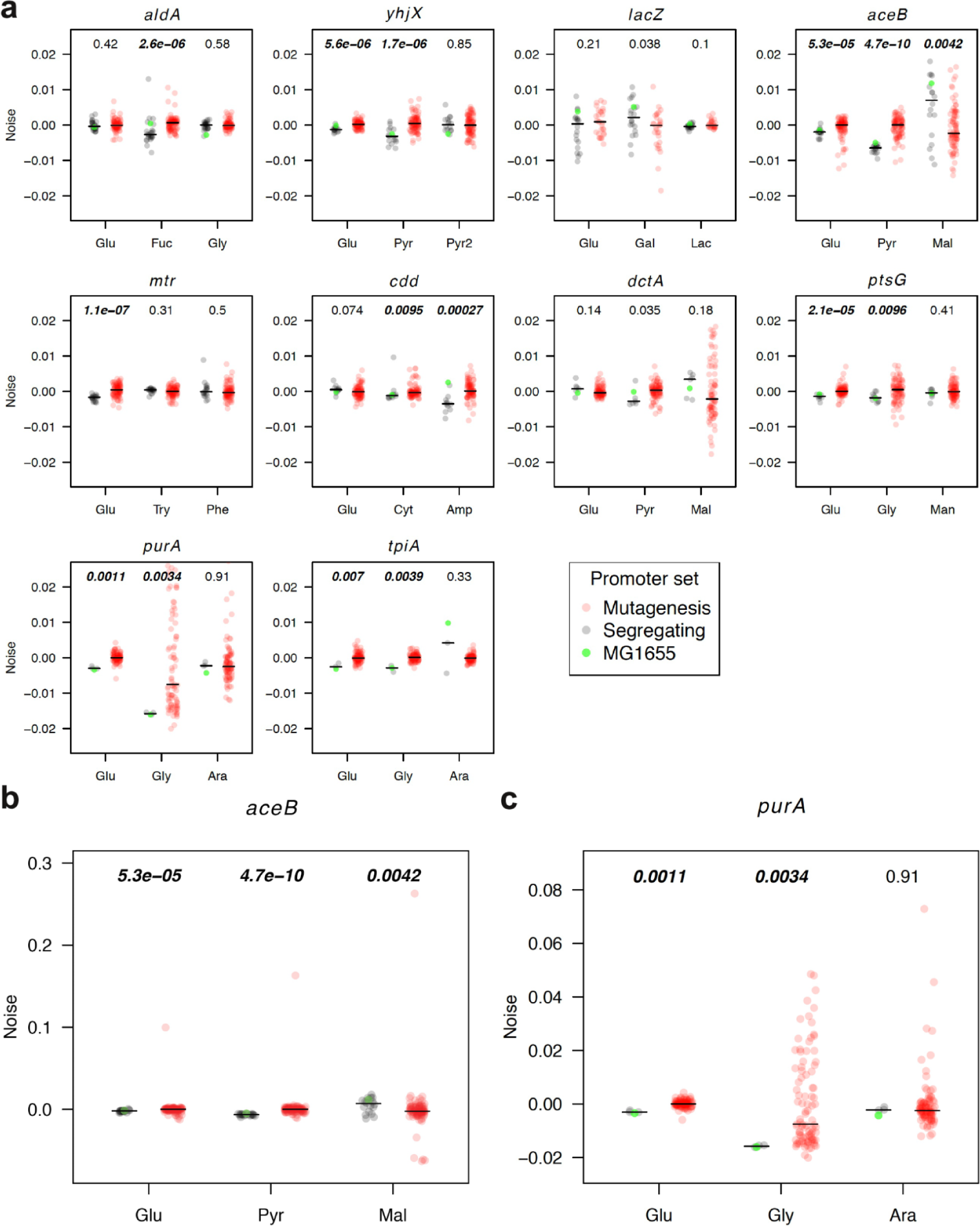
Selection on noise. Segregating promoter variants are selected such that they have low noise in most environments. Here, noise refers to the deviation from a spline fitted of modal population expression vs. the standard deviation in expression divided by the modal population expression (**Supplementary** Figure 4). **a**) Differences in noise between segregating (black) and random (red) variants. The black lines indicate the median noise value for each group. We tested for differences in noise between segregating and random variants using the Wilcoxon rank-sum test; the resulting p-values are indicated above each pair. The numbers in bold indicate significant p-values after the Bonferroni correction for multiple comparisons within the promoter. The specific environment in which the promoters were tested is indicated on the x-axis (**Table 2**). The panels are arranged in decreasing order of segregating genetic variation (PSS). The MG1655 variant of the *mtr* promoter was omitted from calculation due to a SNP in GFP (**Supplementary Note**). **b**) and **c**) show noise deviations for the *aceB* and *purA* promoters using a wider scale.

However, segregating promoter variants did not exhibit lower noise in all cases. Specifically, segregating *aceB* variants exhibited higher noise when grown in L-malic acid (median noise deviation 7e-03 for segregating variants compared to -2e-03 for random variants, p = 0.004). Notably, in the two other environments that we tested, pyruvic acid and glucose, segregating *aceB* variants exhibited lower noise. In 13 cases out of 30 we did not detect a significant difference in noise between the two groups. In these cases, we expect that the differences are below our limit of detection, or that random mutations are just as likely to result in increased noise as they are in decreased noise. Finally, we note that we have measured noise in only a very limited number of environments; it is possible that in other environments, such as those with high stress, the majority of promoters would exhibit higher noise.

### Segregating polymorphisms are enriched for mutations with small phenotypic effects across multiple expression phenotypes

So far we have focused on how natural selection affects individual expression phenotypes: expression level, phenotypic plasticity, and noise. However, in natural environments all these aspects are simultaneously under selection. While a particular random variant may exhibit behaviour comparable to the segregating variants for a single phenotype, this may not be the case across all the phenotypes. To compare the behaviours of segregating and random variants across all the expression phenotypes, we first made the assumption that regulatory phenotypes are under stabilizing selection, and thus that the mean phenotype of the segregating promoters is close to the optimum. This is clearly supported by the above results, in particular that segregating mutations are enriched for mutations of small effect. Under this assumption, regulatory phenotypes that deviate from this mean are less fit. Thus, we calculated the mean and standard deviation of each phenotype for all segregating variants. We then used these values to calculate a z-score for all variants and phenotypes. We converted these z-scores into absolute values to focus only on the relative difference of the phenotype of the variant, rather than the directionality. Finally, given that we have little information on which of these phenotypes is under stronger selection, we weighed all phenotypes equally, and calculated the sum of absolute values of the z-scores. This metric is indicative of how individual variants differ in their overall phenotype set from the phenotype of the average segregating variant. We found a correlation between the z-scores of different expression levels, as well as between expression levels and plasticity. For expression levels we thus used z-scores from just one out of the three environments. This eliminated the non-independence of individual z-scores among expression levels. However, some non-independence between the z-scores of expression level and plasticity remained (**Methods**).

Using this metric, we found that in all cases except for *lacZ*, random variants exhibited significantly higher cumulative z-scores than did segregating variants (**Fig. 8**). Surprisingly, some random promoter variants (e.g. *purA*) exhibited cumulative z-scores over 100, indicating that their phenotypes were more than 100 standard deviations outside of the segregating variants. In the case of both *purA* and *tpiA*, all random variants had cumulative z-scores higher than the highest z-score of any segregating variant. These results highlight that although for individual phenotypes there are not necessarily strong consistent differences between all segregating and all random variants, considering all phenotypes together clearly illustrates that random mutations generally have much larger effects on regulatory phenotypes than do segregating mutations, and that such mutations are effectively filtered by selection.

**Figure 8:**
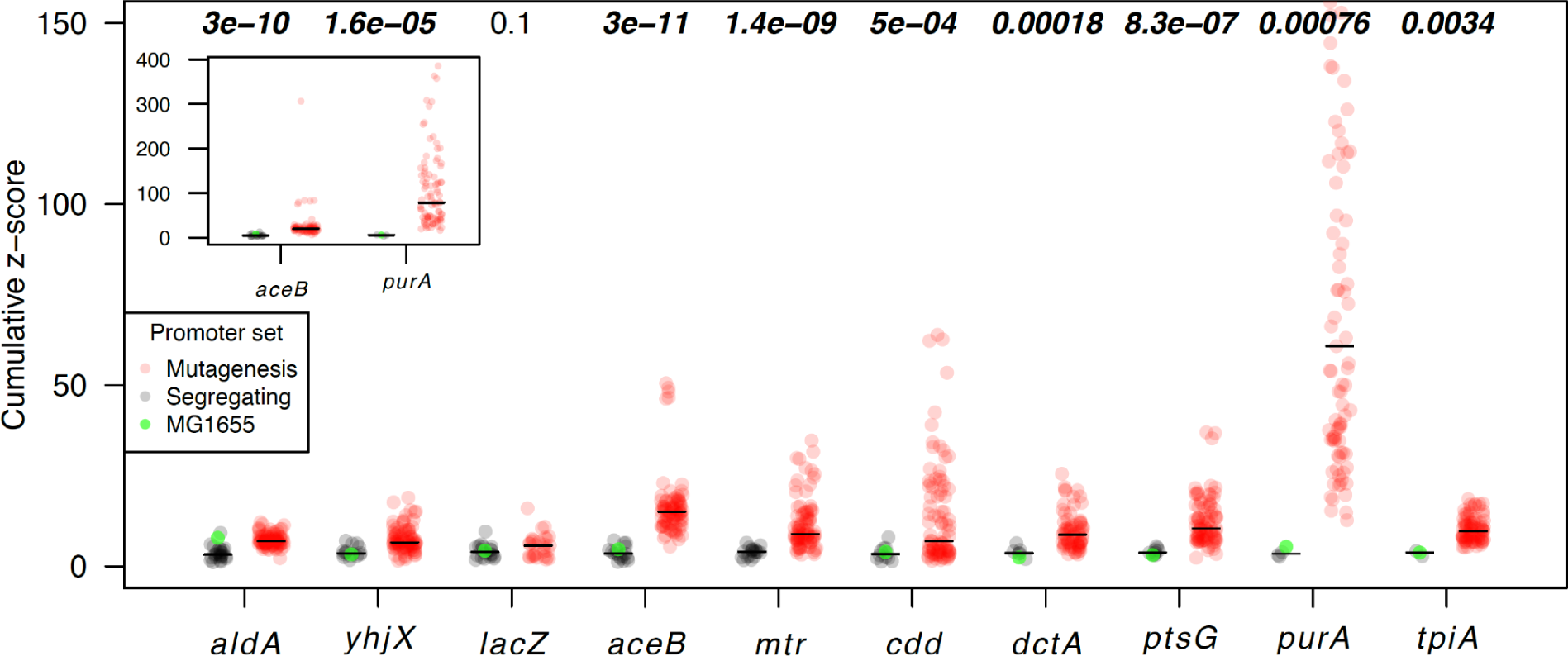
Overall selection pressure. Comparison of cumulative z-scores of all measured phenotypes (expression level, plasticity, and noise) between segregating and random variants. The numbers above each pair of segregating and random variants indicate the p-values for a Wilcoxon rank-sum test to test for differences between the two groups. The numbers in bold indicate significant p-values. The horizontal black lines indicate the median values of cumulative z-scores of each group. The promoters are arranged in decreasing order of segregating genetic variation (PSS). The MG1655 variant of the *mtr* promoter was omitted from calculation due to a SNP in GFP (**Supplementary Note**). The inset shows the full scale of cumulative z-scores on the y-axis for comparison of high values in *aceB* and *purA* promoters.

## Discussion

The effect of selection on gene expression phenotypes has not been well-studied, despite gene expression phenotypes having critical effects on cellular physiology and behaviour. To gain insight into the action of selection on gene expression, here we have compared the phenotypes of segregating variants for ten promoters having different levels of sequence conservation. To understand how selection has acted, we quantified three important phenotypes: modal population expression level, phenotypic plasticity, and noise.

By comparing the variation in expression levels to the level of segregating genetic variation, we established that promoters with higher levels of genetic variation also have higher levels of phenotypic variation (**Fig. 3** and **Supplementary Figure 2**). This suggests that the decreased genetic variation observed in some promoters is due to stronger stabilizing selection.

To more directly measure the action of selection, we created a large number of random variants for each of the ten promoters.To create these we used MG1655 variants, which proved to be representative of segregating variants in the majority of cases. As the random variants were produced via PCR random mutagenesis, they had never been subject to selection. We first quantified the effects of these random unselected mutations on expression level. This comparison highlighted the necessity of using multiple environments to understand mutational effects, as we frequently observed environment-specific effects on expression (**Fig. 4**). Only a small number of studies have examined expression across multiple environments to understand promoter evolution (Duveau et al., 2017; Schaerli et al., 2018; Urchueguía et al., 2019); many more have used only a single environment (Duveau et al., 2018; Hodgins-Davis et al., 2019; Metzger et al., 2015; Schmiedel et al., 2019; Silander et al., 2012). However, bacteria often inhabit multiple different environments.

By comparing expression levels from segregating and random promoter variants, we showed that natural selection has acted such that segregating variants tend to have small-effect mutations. However, the strength of the observed differences between the two groups varies among both promoters and environments. In general, promoters are more robust to mutations affecting expression level when expression is low or completely off (**Fig. 5**, *yhjX*, *mtr* or *dctA* in glucose). More specifically, we observed few strongly repressed promoters exhibiting increased expression due to random mutations.

Using a similar strategy to that above, we compared the phenotypic plasticity of segregating promoter variants and random promoter variants. Overall, we found that five out of ten random variants exhibited lower plasticity. In these cases, there may be stabilising selection on plasticity in natural populations. However, given that random variants generally exhibited lower plasticity, this leaves open the possibility that there is strong directional selection for regulation to be as plastic as possible, but that higher levels of plasticity are difficult to achieve as there is not sufficient genetic variation.

The random variants of *yhjX, lacZ*, *purA*, and *tpiA* did not exhibit different levels of plasticity than segregating variants. Interestingly, three out of four of these promoters are regulated solely via activation (*yhjX*) or repression (*purA* and *tpiA*). This contrasts with all the promoters for which random variants exhibited lower plasticity than segregating variants - all five are regulated by both activation and repression. It is possible that plastic responses from promoters with this simplified architecture are more robust to random mutations, while regulation from promoters having both repressors and activators is easily disrupted.

For *dctA*, we observed significantly higher plasticity for random variants compared to segregating variants in one pair of environments (**Fig. 6e**, pyruvic acid vs. L-malic acid, p = 0.009). This result suggests that the *dctA* promoter is under selection to maintain the same expression level in both environments, and surprisingly, that this regulatory robustness is easily destroyed via very few mutations (**Fig. 6d**) - most random variants differ from the MG1655 variant by only one or two mutations.

Finally, we have shown that cell-to-cell expression variability, or noise, is often subject to strong selection - as has been shown previously (Hornung et al., 2012; Metzger et al., 2015; Rossi et al., 2019; Schmiedel et al., 2019; Silander et al., 2012; Urchueguía et al., 2019). Importantly, the evidence we present here suggests that selection acts directly to decrease noise, rather than this being a correlated response to other selection pressures. In addition, we found that even single mutations have subtle but detectable effects on noise.

It is very possible that in cases in which we did not find significant changes in a phenotype between segregating and random variants it is possible that those promoters are not optimized for certain phenotypes in the environments considered. This might be true especially in cases when the bacteria do not encounter such environments very often.

We did not observe any strong correlations between the extent of segregating variation (**Fig. 1**) and the phenotypic consequences of *de novo* random mutations. For example, there was no evidence that random mutations generally had smaller phenotypic effects in highly polymorphic promoters. There are two possible explanations for this pattern. First, although our assays of expression phenotypes can detect differences between random and segregating mutations, there may be more subtle differences in mutational effects that we cannot detect. For example, there may be mutations with small effects that are detected by selection but not in this experimental context. The second possibility is that *de novo* mutations do have similar effects in highly polymorphic and conserved promoters, but selection is not as strong in highly polymorphic promoters, allowing through more small-effect mutations. This is supported by the correlation we observed between genetic variation and phenotypic variation in expression levels (**Fig. 3** and **Supplementary Figure 2**). To confirm this hypothesis fitness assays are required. However, previous work suggests that fitness differences can be difficult to detect even when expression levels change by more than ten-fold (Keren et al., 2016). In the majority of results here, random mutations have less than two-fold effects on expression level. Thus, the approach we take here, comparing the phenotypes of segregating and random variants, may be a more powerful method to infer the action of selection than assaying the effects of random mutations on fitness relative to a single wild type.

Finally, we showed that taking into account all the regulatory behaviours we have quantified here - expression level, phenotypic plasticity, and noise - that selection has acted such that segregating SNPs have only minimal phenotypic effects. This conclusion is made more compelling through the fact that the sequences of segregating promoter variants differ from each other by, on average, more than three-fold compared to the sequences of random variants. Thus, there are many single mutations that have large phenotypic effects - yet none of these are segregating.

## Methods

### Promoter selection

We downloaded a database of transcription start sites (TSSs) driven by the σ^70^ factor from RegulonDB (Santos-Zavaleta et al., 2019). From this database, we filtered out TSSs denoted as “weak” in RegulonDB, or those outside of intergenic regions (IGRs). We also removed duplicated IGRs if multiple TSSs were present in the same IGR with the same orientation. This filtering resulted in 605 IGRs, each having one or more TSS, that were used for further analysis. Based on the TSS annotations, the IGRs together with 100 bp of flanking upstream and downstream sequences were extracted from a reference MG1655 genome using GenBank file annotation (Blattner et al., 1997). Throughout, we refer to IGRs together with the flanking sequences as “promoters”. The genomes of 135 environmental *E. coli* strains (Breckell & Silander, 2020; Ishii et al., 2006; Sakoparnig et al., 2021) were then blasted against the promoter sequences extracted from MG1655 with e-value cut-off set to 10^-10^; Blast version 2.7.1 (Altschul et al., 1990). Blast hits not more than 100 bp shorter than the MG1655 reference sequence were extracted as segregating promoter variants from each of the 135 environmental *E. coli* strains. All segregating promoter variants of each promoter were then aligned via t-coffee version 11.0.8 with parameter mode procoffee (Notredame et al., 2000). We then calculated the average pairwise identity (API; t-coffee) and proportion of segregating sites (PSS) for each promoter and IGR (excluding the 100 bp flanking sequences).

We also extracted open reading frames (ORFs) directly downstream of each of the 605 filtered promoters from the MG1655 reference genome (GenBank annotation). The same workflow for extracting segregating promoter variants was followed when extracting segregating ORF variants from the 135 strains of *E. coli* (changing the t-coffee parameter setting -type=dna). No extra flanking regions were included in the ORFs. Out of the 605 promoters originally identified in the MG1655 genome, 429 were found in at least 130 of the 135 environmental *E. coli* isolates together with their downstream ORFs. For all further analyses and tests we used only these 429 promoters, IGRs, and ORFs.

To check whether any functional classes of ORFs were enriched for promoters with IGRs that have high or low sequence variation, we grouped the IGRs into the major categories defined by MultiFun (Serres & Riley, 2000) according to their downstream ORF. IGR variation (PSS and API) from each group was then compared to the rest of the groups together using a Wilcoxon rank-sum test. We also tested the correlation in sequence variation (PSS and API) between IGRs only and promoters (IGRs with 100 bp of upstream and downstream sequences flanking each IGR) using Spearman’s correlation test. All the scripts used with the workflow described above can be found in the **Supplementary Information**.

From the resulting promoter dataset we selected ten promoters exhibiting various levels of sequence variation. We aimed at selecting promoters with obvious and easy differential regulation, i.e, either with known environments in which differential expression occurs or those whose annotated transcription factors suggested possible environments for differential expression. The selected promoters - *aldA*, *yhjX*, *lacZ*, *aceB*, *mtr*, *cdd*, *dctA*,*ptsG*, *purA*, and *tpiA* - differ more than ten-fold in their levels of genetic variation, while they all control expression of genes that are involved in bacterial metabolism or transport of material involved in the metabolism.

### Promoter library construction

For each promoter (*aldA*, *yhjX*, *lacZ*, *aceB*, *mtr*, *cdd*, *dctA*,*ptsG*, *purA*, and *tpiA*) we created two libraries, one with segregating promoter variants and a second with randomly mutated promoter variants. The two promoter libraries were aliquoted into separate microplates, except for the *lacZ* promoter, which was aliquoted into a single microplate. Each microplate also contained the MG1655 variant of the promoter, a positive control consisting of the highly active murein lipoprotein (*lpp*) promoter driving GFP expression, and a negative expression control consisting of a promoter-less pUA66 plasmid (Zaslaver et al., 2006). We describe the construction of each library type separately below.

### Variants segregating in *E. coli* population

Both the vector pUA66 backbone (a low-copy number plasmid with a SC101 ori, a strong RBS, and GFPmut2) and segregating promoter variants were PCR amplified using Phusion High-Fidelity DNA polymerase with HF buffer (New England Biolabs). For promoter PCR amplification, 5μl of pooled DNA from strains with different variants of each promoter was used as a DNA template. The primers for promoter amplification contained 17 nucleotide overhangs which were homologous to the ends of the vector backbone for subsequent DNA assembly. All primers used in this study are listed in **Supplementary Table 1**.

For vector PCR amplification, 0.5 ng of plasmid DNA with pUA66 backbone served as a template. After confirming a successful PCR amplification of the products on 1% agarose gel, the template DNA was digested by DpnI from the remaining reaction volume (Li et al., 2011). The reactions were then column-purified and we assembled the PCR amplified vector and promoters using Gibson assembly (Gibson et al., 2009) with NEBuilder® HiFi DNA Assembly Master Mix (New England Biolabs). The assembly mix was then electroporated directly into the electrocompetent MG1655 strain. Transformed colonies which grew on LB agar plates with 50μg/ml Kanamycin were picked for Sanger sequencing across the insert in the vector backbone and stored as glycerol stocks. Clones with confirmed promoter variants matching the segregating ones were then grown in liquid LB with Kanamycin and used to create 96 well microplate glycerol stock libraries.

### Random variants from PCR mutagenesis

We amplified the pUA66 backbone, DpnI treated, and column-purified it the same way as described for segregating promoter variants above. We produced the promoter inserts by performing error-prone PCR using the GeneMorph II Random Mutagenesis Kit (Agilent Technologies). We used the plasmid constructs with MG1655 promoter variants cloned into them as template DNA for the error-prone PCR aiming to achieve approximately 1.5 SNPs per promoter sequence. We used the same primers as for the segregating promoter variants (**Supplementary Table 1**). Each reaction with randomly mutated promoter variants was column-purified before Gibson assembly with the NEBuilder® HiFi DNA Assembly Master Mix (New England Biolabs). Each assembly mix was then electroporated into the MG1655 strain and colonies that grew on LB with Kanamycin were picked for Sanger sequencing and stored as glycerol stocks. Clones which had none or more than three SNPs in the cloned promoter insert were excluded as well as those with SNPs detected in the vector backbone (rare occasion). The rest of the strains were then re-grown in liquid LB with Kanamycin in 96 deep-well microplates overnight and 96 well microplate glycerol stock libraries were prepared from them.

### Bacterial clones and environments

We performed all the promoter activity assays in the MG1655 genetic background of *E. coli*. All the clones had a pUA66 vector (Zaslaver et al., 2006) differing by promoter variants cloned upstream of the GFPmut2 gene (for details see *Promoter library construction* in **Methods**). We also included positive and negative expression controls which consisted of the highly active plpp::GFPmut2 strain from the Alon Zaslaver library collection and promoter-less pUA66 vector, respectively (Zaslaver et al., 2006).

The assays were performed in 96 well microplates in M9 minimal media (Sigma-Aldrich) supplemented with MgSO_4_, CaCl_2_, a carbon source, and any additional reagents needed to induce or repress expression from a particular promoter (**Table 2**). We grew all the clones in the presence of 50µg/ml Kanamycin to prevent plasmid loss.

Before introducing the clones into the assay media we first inoculated them into 0.5ml of M9 minimal media with 0.4% glucose and Kanamycin (M9 glucose) from a 96 well microplate glycerol stock using a 96-well pin replicator (Enzyscreen B.V.). We then incubated these microplates overnight at 37°C with shaking to revive the cells. After overnight growth in M9 glucose, we inoculated each grown clone into 0.5ml of one of the assay media in a 96 deep-well microplate using the pin replicator and incubated them for 24h at 37°C with shaking. After this second incubation we inoculated the clones into the same fresh assay media they grew in for 24h, but into three separate 96 deep-well microplates (to obtain triplicates for each clone). These cultures were incubated until they reached an exponential phase (**Table 2**).

### Flow cytometry analysis

Once the cells reached exponential growth (**Table 2**) in the appropriate assay media, we diluted them into 1x PBS with ∼2.5% formaldehyde and kept them on ice until performing flow cytometry analysis the same day. We performed the flow cytometry on a BD FACSCanto II machine using BD FACSDiva software version 6.1.3. We obtained the GFP fluorescence data through the 488 nm laser and a 513/17 nm bandpass filter. We set the number of events to record from each well to 20,000. We exported the acquired data from FACSDiva software into Flow Cytometry Standard files, and performed all cell gating and fluorescence analysis using custom R scripts (flowCore package version 2.0.1; see **Supplementary Information**). We gated cells based on their maximal kernel density of forward and side scatter values, using the ellipsoidGate function from the flowCore package, and keeping about ⅓ of all events. The modal population expression was calculated as the mean of the three maximal kernel density values from the GFP fluorescence signal of three replicates. The coefficient of variation (CV) was calculated separately for each of the three biological replicates (standard deviation divided by the modal population expression), and the mean of these values was used as the CV of the promoter. Replicates with fewer than 2,500 and 5,000 recorded events were excluded from the calculation of modal population expression and coefficient of variation, respectively. We found that when fewer than 5,000 events were collected, a larger number of these were likely to be outlier events (e.g. machine noise), which affected the calculations of variance. When comparing segregating and random variants of the same promoter that came from two separate microplates, we obtained an offset for both modal population expression and coefficient of variation to minimise plate-effects. We calculated these offsets as the mean of the differences between the three controls present in each microplate, i.e., the MG1655 promoter variant, plpp::GFPmut2 (positive control), and pUA66 (promoterless negative control). All figures contain modal expression and coefficient of variation values that are corrected using these offsets. All scripts using the workflow described here, including the original data files can be found in the **Supplementary Information**.

### Testing the correlation between segregating genotypic and phenotypic variation

To compare sequence promoter variation with variation in expression levels among segregating variants we used the promoter sequences (IGRs with 100 bp flanking regions) of the ten selected promoters (*aldA*, *yhjX*, *lacZ*, *aceB*, *mtr*, *cdd*, *dctA*,*ptsG*, *purA*, and *tpiA*). We measured modal population expression level in triplicates from each segregating promoter variant in three different environments (**Table 2**), and used the mean of these triplicate measures. We then calculated the standard deviation of modal population expression levels from all segregating variants of each promoter in each environment as a measure of phenotypic variation. For each promoter we thus had three values of segregating phenotypic variation (one for each environment). The MG1655 variant of the *mtr* promoter was excluded from the calculations due to a SNP in GFP causing lower fluorescence (**Supplementary Note**). We then used Spearman’s correlation test to calculate the correlation between the phenotypic variation and genotypic variation using four metrics: PSS, API, total number of existing segregating variants and number of cloned segregating variants (the code for these calculations can be found in the Supplementary Information).

### Mapping the effects of single SNPs to promoter sequence

When mapping the effect size and direction of randomly introduced SNPs we used modal population expression values only from those random variants which had contained a single SNP relative to the MG1655 promoter variant. Information about the TF binding sites, -10 and -35 elements, and ORFs was taken from the EcoCyc database (Karp et al., 2018), and only the annotations associated with σ^70^ driven TSSs were used. Using the Sanger sequencing results, we identified the location of all SNPs for each random variant (the code for these calculations can be found in the **Supplementary Information**).

### Comparison of phenotypic variation between segregating and random variants

To test for differences in the variation in expression between the segregating and random variants, we calculated the modal population expression values for each promoter variant in both groups and all environments. We then tested for significant differences in variation in modal population expression levels using the Fligner-Killeen test of homogeneity of variances. We included values from the non-mutated MG1655 promoter variants in the sets with the segregating variants, except for the *mtr* promoter due to a SNP in GFP (**Supplementary Note**; the code for these calculations can be found in the **Supplementary Information**).

### Comparing plasticity between segregating and random variants

We calculated the phenotypic plasticity of promoter variants across all three environments by calculating the Euclidean distance of each datapoint in three dimensions to an isocline representing null plasticity. The isocline is defined by equal values in the three dimensional space, i.e., x = y = z. The three dimensions are defined by the modal population expression values in the three environments specific for each promoter. Each datapoint (promoter variant) is thus defined by its expression values in all three environments. The closer a datapoint is to the isocline, the lower the plasticity.

To calculate plasticity in pairs of environments, we used an analogous method for two environments. To test whether natural selection had acted on plasticity (across all three environments and in pairs of environments) we compared plasticity values from all segregating (including MG1655 variants, except the one from *mtr* promoter - **Supplementary Note**) and all random variants for each of the ten promoters using the Wilcoxon rank-sum test (the code for these calculations can be found in the **Supplementary Information**).

### Comparing noise between segregating and random variants

To determine the noise within an isogenic cell population of each promoter variant, we first excluded fluorescence values from the population that were lower or higher than three standard deviations from the modal population expression level. Then we calculated the coefficient of variation from this isogenic cell population as the standard deviation of the fluorescence divided by the modal population expression level. We next fitted a cubic smoothing spline (smoothing parameter lambda = 0.01) to the modal population expression vs. the coefficient of variation values, using all (segregating and random) promoter variants. We determined the noise levels as the difference in measured coefficient of variation from the coefficient of variation predicted from the fitted spline. We then compared the noise values between segregating and random variants using Wilcoxon rank-sum test to determine whether selection had acted on noise (the code for these calculations can be found in the **Supplementary Information**). The MG1655 variant of the *mtr* promoter was excluded from the calculations due to a SNP in GFP causing lower fluorescence (**Supplementary Note**).

### Comparing overall promoter activity between segregating and random variants

We calculated the mean and standard deviation for each promoter and phenotype (modal population expression in all three environments, plasticity across all three environments and transcriptional noise deviation for each of the three environments) using just segregating variants. We then used these mean and standard deviation values of segregating variants to calculate z-score for each individual promoter variant (both segregating and random) to determine how much it differs in a particular phenotype from an average segregating variant. To focus on the relative size of the change, rather than on directionality, we converted all z-scores into absolute values. Giving all the phenotypes an equal weighting, we summed the z-scores for each variant to measure its deviation from an average segregating variant in all the phenotypes together. Due to a strong correlation between the z-scores for expression levels in the three environments, we used only expression level z-scores from a single environment. This environment was chosen so that the median expression from the segregating variants was higher than 2.5 to remove environments causing promoter repression. If more than one environment remained, we used the one least correlated with plasticity. This was done to minimize non-independence of individual z-scores when calculating cumulative z-scores. We then checked for statistically significant differences between the cumulative z-scores of segregating (including MG1655 variants, except the one from *mtr* promoter - **Supplementary Note**) and random variants using Wilcoxon rank-sum test (the code for these calculations can be found in the **Supplementary Information**).

## Acknowledgements

We thank Tim Cooper, Andrea Sajuthi, and Nikki Freed for valuable comments on the final draft of this manuscript. We are also grateful to Stella Pearless and Bhargava Morampalli for sequencing several plasmid constructs. This work was supported by a Marsden Grant (grant MAU1703) awarded to OKS. The funder had no role in study design, data collection and interpretation, or the decision to submit the work for publication.

## Author contributions

MV and OKS conceived the project and designed the experiments and analyses. OKS supervised the project. MV performed all experiments and all analyses. MV wrote the paper with contributions from OKS.

## Conflict of interests

The authors declare that there are no conflicts of interests.

## Data availability

The original data files and scripts with data analyses that support the findings of this study are available in the **Supplementary Information** of this study.

## Supplementary Information

All scripts with access to original data files can be found at https://github.com/marketavlkova/2021_PromoterEvolution, the Supplementary Files can be accessed through Figshare DOI: https://doi.org/10.6084/m9.figshare.c.5517228.v2.

## Supplementary Note

We found that the MG1655 *mtr* variant is expressed at lower levels than all other *mtr* segregating variants when grown in glucose with phenylalanine. In addition, the promoter variants generated via random mutagenesis of the MG1655 variant exhibited expression levels similar to the segregating variants rather than MG1655. We thus sequenced the plasmid containing the MG1655 promoter variant, and found that there was a single non-synonymous mutation in GFP at position 521 changing the codon GGA (G) to GAA (E). A small fraction of other constructs may also contain a mutation outside of the promoter region, but we expect that the large sample sizes we use will negate the effect of this small fraction on our conclusions.

**Supplementary Figure 1:**
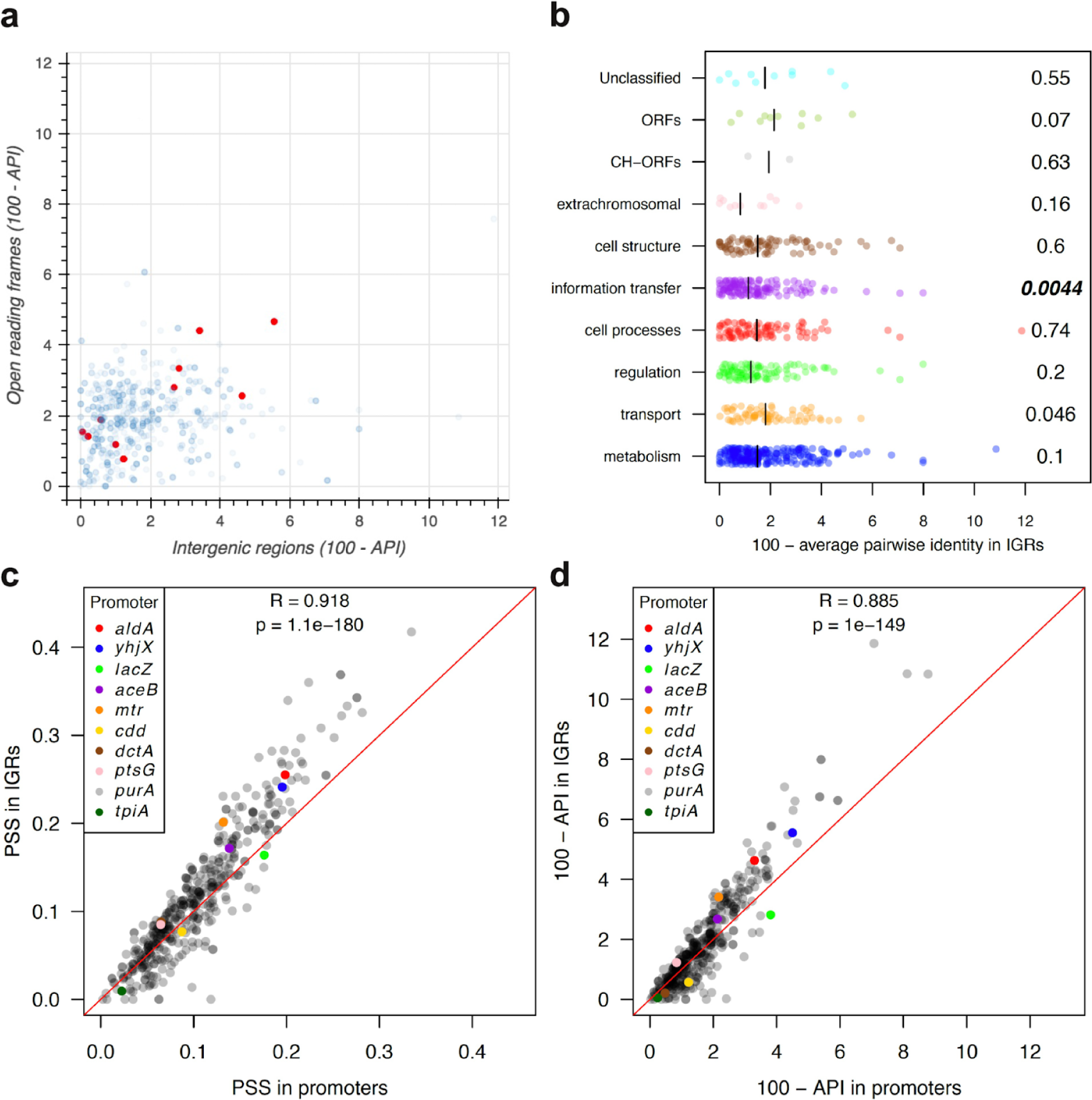
Polymorphisms in intergenic regions (IGRs) and open reading frames (ORFs) across 135 environmental isolates of *E. coli* and MG1655. a) The inverted average pairwise identity (100 - API) for IGRs and downstream ORFs varies by more than an order of magnitude. Each blue dot indicates an inverted API for an IGR-ORF pair when the IGR contains a transcriptional start site for the ORF. In red are IGRs that we selected for further study due to their different levels of sequence variation (*aldA*, *yhjX*, *lacZ, aceB*, *mtr, cdd*, *dctA*, *ptsG*, *purA*, and *tpiA*; see Table 1). b) The inverted API for IGRs differs little among different functional groups, as classified by the downstream ORF. CH denotes conserved-hypothetical. Unclassified, ORFs, and CH-ORFs all represent groups of ORFs with very limited information on function (Serres & Riley, 2000). Numbers next to each function group represent the p-value obtained via the Wilcoxon rank-sum test to test for differences in API in each group from all other inverted API values for IGRs outside of the group. Bold indicates significant p-values after the Bonferroni correction for multiple comparisons among functional groups. The black lines indicate the mean inverted API values of each functional group. c) and d) Correlation in sequence variation between IGRs and promoters (IGRs with 100 bp of flanking ORF regions). c shows the proportion of segregating sites (PSS) and d displays the inverted average pairwise identity (100 - API).

**Supplementary Figure 2:**
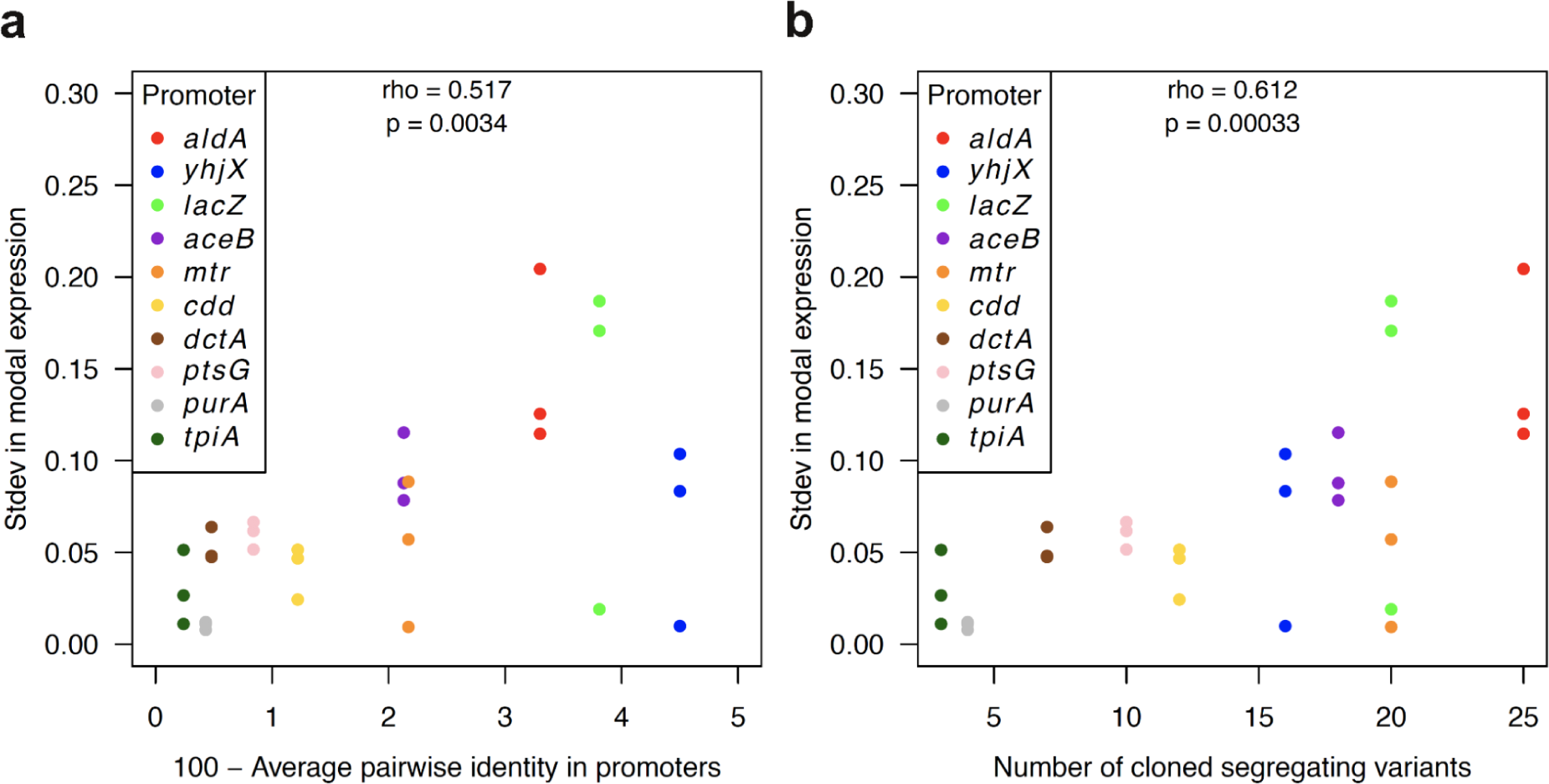
Segregating genetic variation in promoters correlates with variation in expression levels. a) and b) Standard deviations in modal expression levels from segregating variants are correlated with the genetic variation of the promoter (IGR with 100 bp flanking regions). Panel a shows the correlation with API and panel b shows the correlation with the number of segregating promoter variants cloned and used for phenotypic assay. For each promoter, the standard deviation of modal population expression was measured in three environments (three dots per promoter, Table 2). The rho and p-values were calculated using Spearman’s correlation test.

**Supplementary Figure 3:**
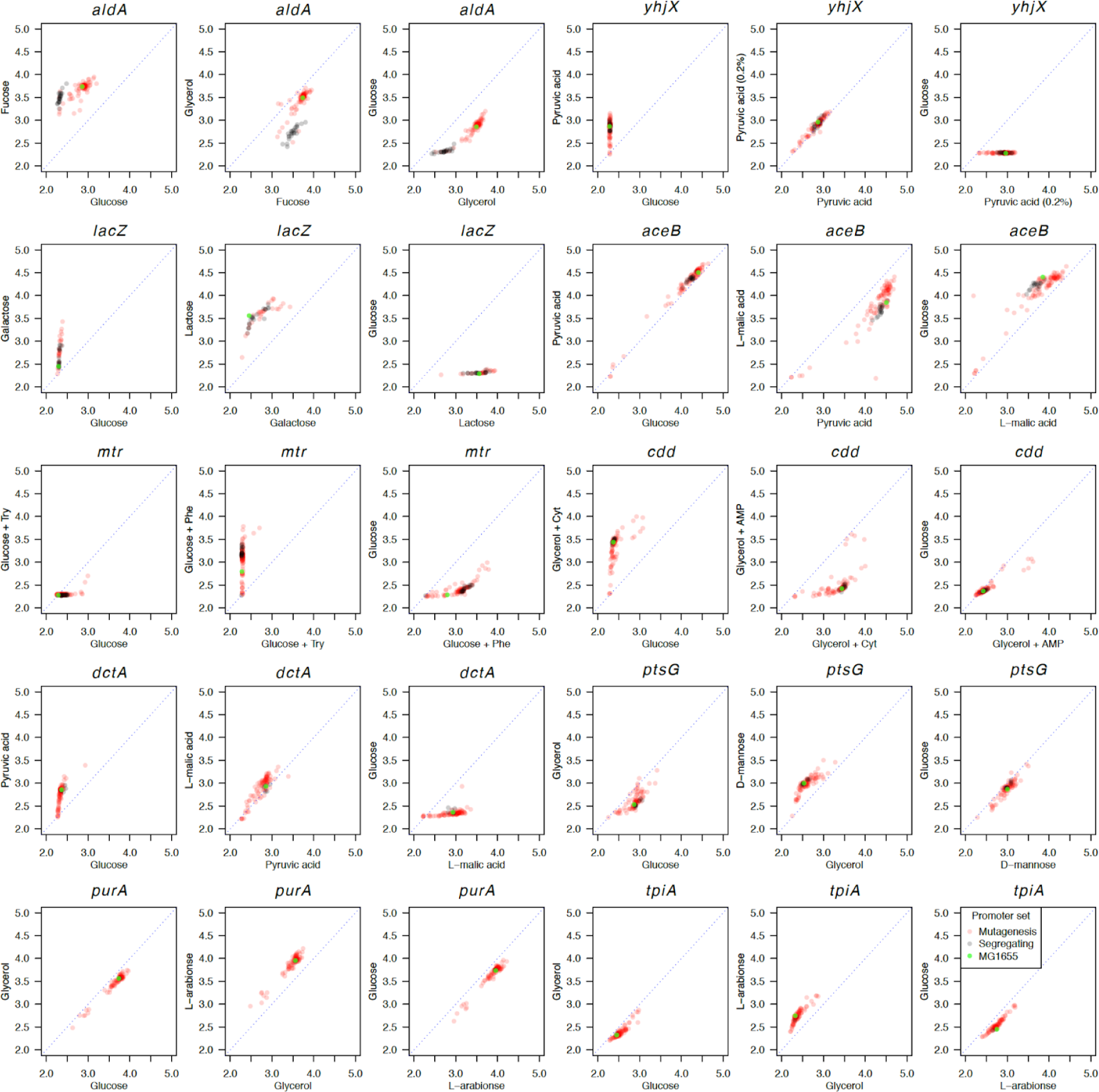
Comparison of expression levels from promoters in pairs of environments. All x and y axes represent the modal population expression level in the particular environment written on the axis label. The blue dotted lines indicate equal expression levels in both environments, i.e, no phenotypic plasticity. The further from the line a promoter is, the higher the absolute difference is in expression from the promoter between the two environments (i.e., the higher its phenotypic plasticity).

**Supplementary Figure 4:**
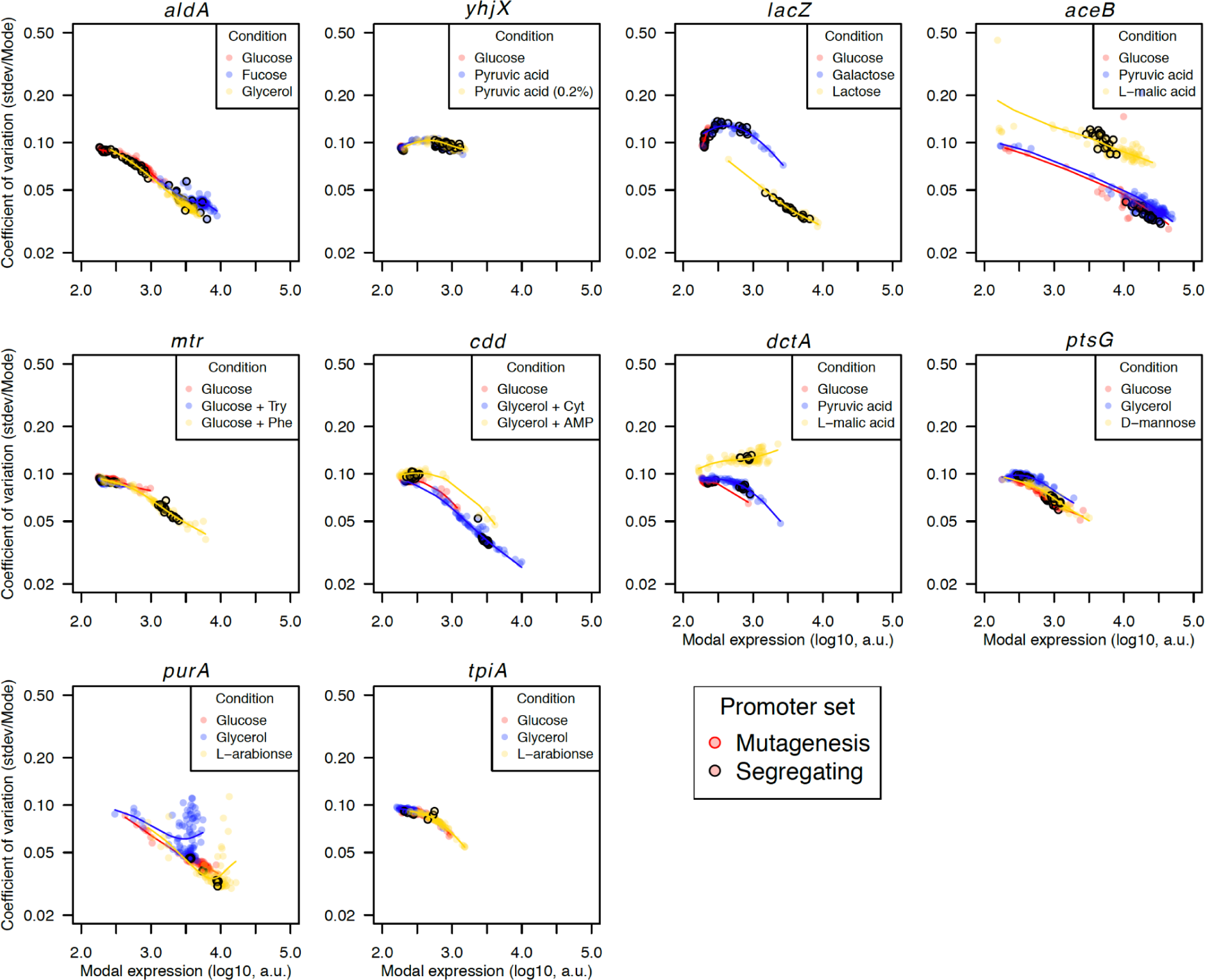
Fits of smoothing splines to modal population expression and coefficient of variation. A smoothing spline was fitted to all variants (segregating and mutagenized) in each environment. The term “noise” is used for the vertical deviation of each variant, i.e., deviation in the coefficient of variation from the fitted spline. The coefficient of variation is calculated as a standard deviation of log transformed expression levels (stdev) divided by modal population expression level (Mode).

**Supplementary Figure 5:**
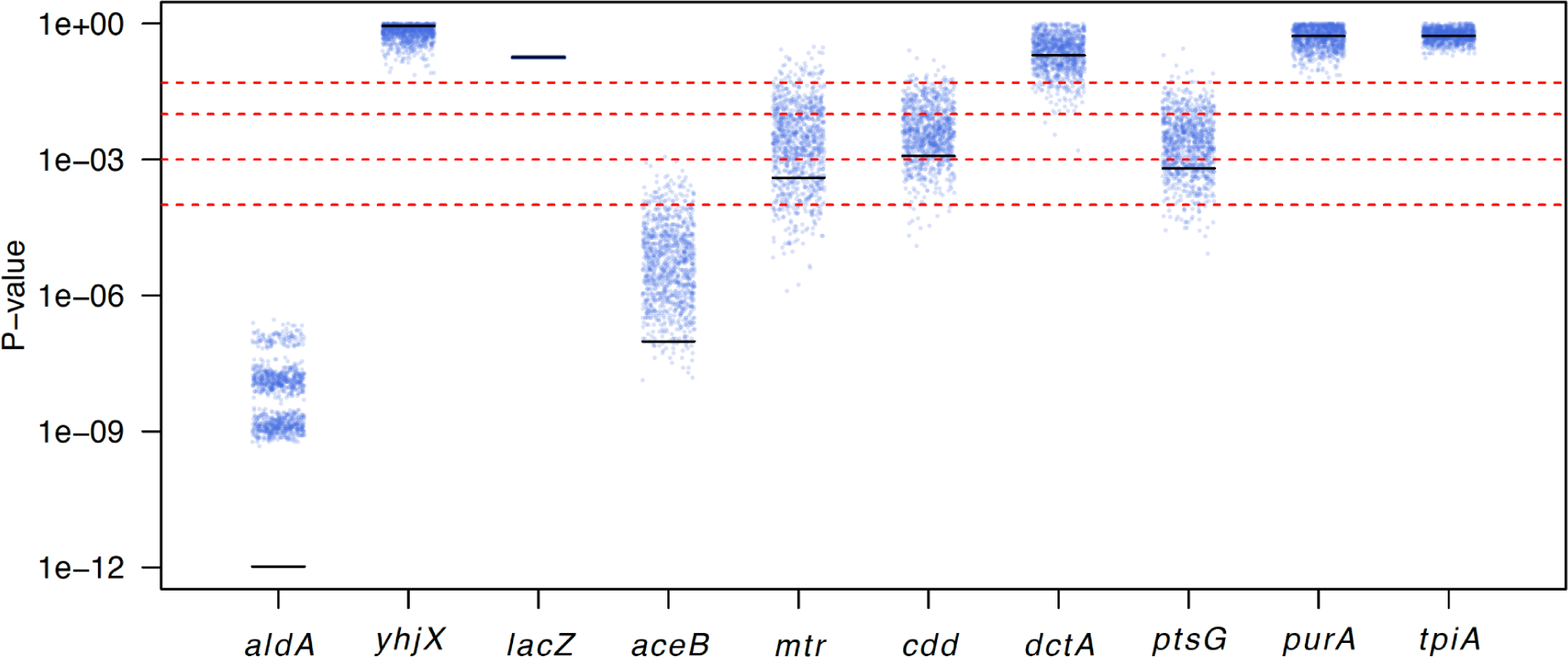
The lower sample sizes of random variants can increase p-values by two orders of magnitude. To test whether the not-significant p-value for *lacZ* promoter in plasticity (as in Fig. 6c) could be due to the lower number of random variants generated, we subsampled random variants of all other promoters to match the number of random *lacZ* variants (29 in total). We repeated this subsampling 1000-times, and for each iteration we compared the 29 subsampled random variants to all the segregating variants using Wilcoxon rank-sum test. This shows that the lower sample size for *lacZ* random variants likely decreased our power to find differences between segregating and random variants. The resulting p-values are plotted here. The horizontal red lines indicate p-values of 0.05, 0.01, 0.001, and 0.0001. The black lines indicate the p-values without subsampling (as in Fig. 6c).

## Notes

### Competing Interest Statement

The authors have declared no competing interest.

